# Adaptive diversification in the cellular circadian behavior of *Arabidopsis* leaf- and root-derived cells

**DOI:** 10.1101/2021.06.10.447857

**Authors:** Shunji Nakamura, Tokitaka Oyama

## Abstract

The plant circadian system is based on self-sustained cellular oscillations and is utilized to adapt to daily and seasonal environmental changes. The cellular circadian clocks in the above- and belowground plant organs are subjected to diverse local environments. Individual cellular clocks are affected by other cells/tissues in plants, and the intrinsic properties of cellular clocks remain to be elucidated. In this study, we showed the circadian properties of leaf- and root-derived cells of a *CCA1::LUC Arabidopsis* transgenic plant and demonstrated that the cells in total isolation from other cells harbor a genuine circadian clock. Quantitative and statistical analyses for individual cellular bioluminescence rhythms revealed a difference in amplitude and precision of light/dark entrainment between the two cell-types, suggesting that leaf-derived cells have a clock with a stronger persistence against fluctuating environments. Circadian systems in the leaves and roots are diversified to adapt to their local environments at the cellular level.

## INTRODUCTION

The circadian clock is a biological system possessed by most organisms that enables them to adapt to daily and seasonal environmental changes. The circadian rhythm satisfies three fundamental properties, self-sustained oscillation with an approximate 24-hour period under constant conditions, temperature-compensation of the period, and entrainability to the day-night cycles through responses to environmental changes, such as light/dark and high/low temperature^1^. Therefore, the circadian system of plants meets the seemingly competing demands, i.e., responsiveness to input signals from environmental cues and the periodicity enabling an invariant circadian cycle under different environments^2–6^ . In addition, circadian oscillation in plants could be perturbed by environmental fluctuations caused by changes in weather patterns as external noise and internal fluctuations of clock components as molecular noise^7–9^. In natural habitats, the degrees of daily changes and high-frequency fluctuations in environmental factors are larger above the ground than in the soil, and the circadian system in the root is likely different from that in the shoot^10–13^.

In plants, the circadian clock is based on self-oscillatory transcription-translation feedback loops of clock genes^14^ . The self-oscillatory gene circuit is associated with light signaling, and its timing is correctly entrained in the day-night cycles^5^. The circadian expression of clock genes influences the temporal regulation of various physiological processes. Hence, the circadian system of plants is based on the cell-autonomous machinery^6,13^. In addition to cell-autonomous basis, the circadian timing in the plant body is coordinated in association with intercellular communication^15^. Exploring the circadian behavior of individual cells without the influence of intercellular communication has been a challenge^15^. In other words, the cellular basis of the circadian system is still unclear.

The circadian behavior of individual cells in plants have been directly observed by using fluorescent and bioluminescent reporter systems^12,13,16–20^. Under constant conditions, desynchronization of cellular rhythms occurs due to heterogeneous and unstable properties of individual cellular rhythms. Even under such asynchronous conditions, it has been suggested that intercellular communication influences the circadian behavior of cells^12,13,20–24^. The use of isolated cells (protoplasts) and suspension-culture cells, which prevents the influence of tissue/organ systems^25–27^, has been adopted for cell-based studies on the circadian rhythms of plant. Cell density affects the circadian behavior of cells, which implies that intercellular communication between isolated shoot apex cells influences the circadian rhythm^12^.

In this study, we performed a single-cell analysis on the circadian behavior of *Arabidopsis* cells separated from other cells. Data obtained during the bioluminescence monitoring of individual cells isolated from transgenic plants carrying a luciferase (*LUC*) fusion gene under the control of the *CIRCADIAN CLOCK ASSOCIATED 1* (*CCA1*, a core clock gene) promoter were subjected to quantitative and statistical analyses. We demonstrated the three fundamental properties, and cell-type specificity of circadian rhythms in leaf- and root-derived cells. Furthermore, we discussed the adaptation of cellular circadian clocks of plants to fluctuating environments.

## RESULTS

### Experimental setup for monitoring the bioluminescence of individual protoplast-derived cells

We established a bioluminescence monitoring system to observe the circadian rhythms of protoplast-derived cells at a single-cell level (Supplementary Fig. 1). As previously reported^27–30^, we prepared protoplasts from the leaves and roots of an *Arabidopsis CCA1::LUC* transgenic plant. Protoplasts of leaves are mainly mesophyll cells^27^ and those of roots are presumedly a heterogeneous population with different cell types^31,32^. Cells derived from the protoplasts of leaves and roots are hereafter referred to as leaf-derived cells and root-derived cells, respectively. Before isolating the protoplasts, the plants were entrained (synchronized) through exposure to two cycles of 12-hour light/12-hour dark (2×LD). We monitored the bioluminescence of the protoplast-derived cells by using a cooled electron-multiplying charge-coupled-device (EM-CCD) camera under a macro zoom microscope. As the standard conditions during bioluminescence monitoring, the protoplast-derived cells were cultured in a liquid medium (W5FBS with 0.1 mM luciferin; 2 × 10^4^ cells/ 4 ml for leaf-derived cells, 5 × 10^4^ cells/4 ml for root-derived cells) under constant dark (DD) conditions at 22°C. We also applied experimental conditions altered in light schedules, temperature, or cell densities.

### Grouping luminous cells for circadian rhythm analysis

Luminous cells under DD condition were categorized into cells with an appropriate circadian rhythm (Group 0) and those with irregular rhythm (Group 1 and 2) (Supplementary Fig. 2). Any bioluminescence time series lacking the first peak within 24 h were categorized under Group 2. The cells in this group failed to initiate the bioluminescence circadian rhythm or to correctly phase the rhythm. Time series including a 35-hour or longer range without a peak time were categorized under Group 1. The cells in this group included those that lost bioluminescence in the monitoring.

Under standard conditions, ∼30% luminous cells isolated from both leaves and roots were those in Group 1 (Table 1). Interestingly, cells with irregular rhythm appeared to retain the circadian periodicity (Table 1). The distributions of peak intervals of cells with irregular rhythm were similar to those of cells with circadian rhythm, although the distributions of cells with irregular rhythm were more deviated than those of cells with circadian rhythm (Table 1 and Supplementary Fig. 3). With respect to the phasing of the bioluminescence rhythm, the first peaks of both leaf- and root-derived cells (all analyzable cells) appeared 18 h after the light-off at a maximum rate, while the timing of the first peak for leaf-derived cells was much more dispersed than that for root-derived cells (Supplementary Fig. 4). The cells in Group 0 showed contrasting phenomena of temporal changes in the synchrony (Supplementary Fig. 4c,d). The leaf-derived cells started with relatively low synchrony and maintained the synchronous state, whereas the root-derived cells started with relatively high synchrony, which steeply reduced. This suggested that the circadian system of leaf-derived cells had lower preciseness for phasing but seemingly higher preciseness for periodicity than that of root-derived ones. Both leaf- and root-derived cells in Groups 1 and 2 lost their synchrony by the end of the monitoring period (Supplementary Fig. 4c,d). To interpret the time-series data for cellular circadian behavior, we show the results for bioluminescence rhythms of cells only in Group 0 (cells with circadian rhythm) in the following sections, unless described otherwise.

**Table 1.**
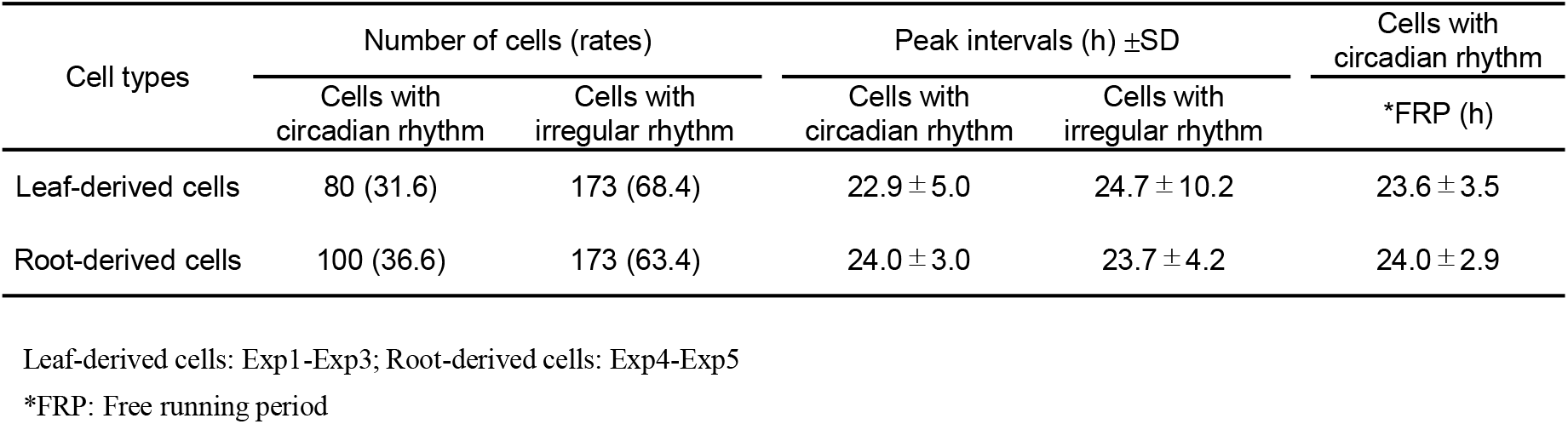
Summary of circadian rhythmicity of protoplast-derived cells under standard conditions

| Cell types | Number of cells (rates) | | Peak intervals (h) ±SD | | Cells with circadian rhythm |
| --- | --- | --- | --- | --- | --- |
|  | Cells with circadian rhythm | Cells with irregular rhythm | Cells with circadian rhythm | Cells with irregular rhythm | *FRP (h) |
| Leaf-derived cells | 80 (31.6) | 173 (68.4) | 22.9 ± 5.0 | 24.7 ± 10.2 | 23.6 ± 3.5 |
| Root-derived cells | 100 (36.6) | 173 (63.4) | 24.0 ± 3.0 | 23.7 ± 4.2 | 24.0 ± 2.9 |

Leaf-derived cells: Exp1-Exp3; Root-derived cells: Exp4-Exp5
\*FRP: Free running period

### Comparison of bioluminescence rhythms of leaf- and root-derived cells under standard conditions

For a cellular process to be defined as a circadian rhythm, it must fulfill three fundamental properties: persistence in constant conditions for an about 24 h period, temperature compensation of the period, and entrainment. As earlier mentioned, we observed a robust oscillation in leaf- and root-derived cells in Group 0 (cells with circadian rhythm) (Figure 1a,b) and circadian periodicity in Groups 1 and 2 (cells with irregular rhythm) (Table 1). With respect to the trends of the luminescence intensity, a large number of leaf-derived cells showed increased luminescence intensity during the monitoring, whereas many root-derived cells had decreased luminescence intensity (Supplementary Fig. 5). Root-derived cells may have a damping oscillator and show a high responsiveness to external stimuli, such as light/dark. The cellular period of a cellular rhythm is defined as the mean of peak intervals of a cellular rhythm. The cellular periods of both leaf- and root-derived cells were widely distributed with a mean value of approximately 24 h, indicating that the cells retained circadian rhythmicity as a whole (Fig. 1c, Table 1and Supplementary Table 1). To obtain information on the cycle-to-cycle variations, we calculated the coefficient of variation (CV = standard deviation/mean) of cellular period for each cellular rhythm. The CV of cellular period of leaf-derived cells was higher than that of root-derived ones, suggesting that bioluminescence circadian rhythms of leaf-derived cells were unstably generated (Fig. 1d and Supplementary Table 1). Taken together, with the seemingly higher preciseness of the periodicity of leaf-derived cells (Supplementary Fig. 4), the instability of peak times may be due to the instability of bioluminescence generation downstream of the clock rather than the instability of the clock itself. Despite a relatively large variation of peak intervals (Supplementary Fig. 3), leaf- and root-derived cells showed self-sustaining property with a periodicity of approximately 24 h.

**Fig. 1.**
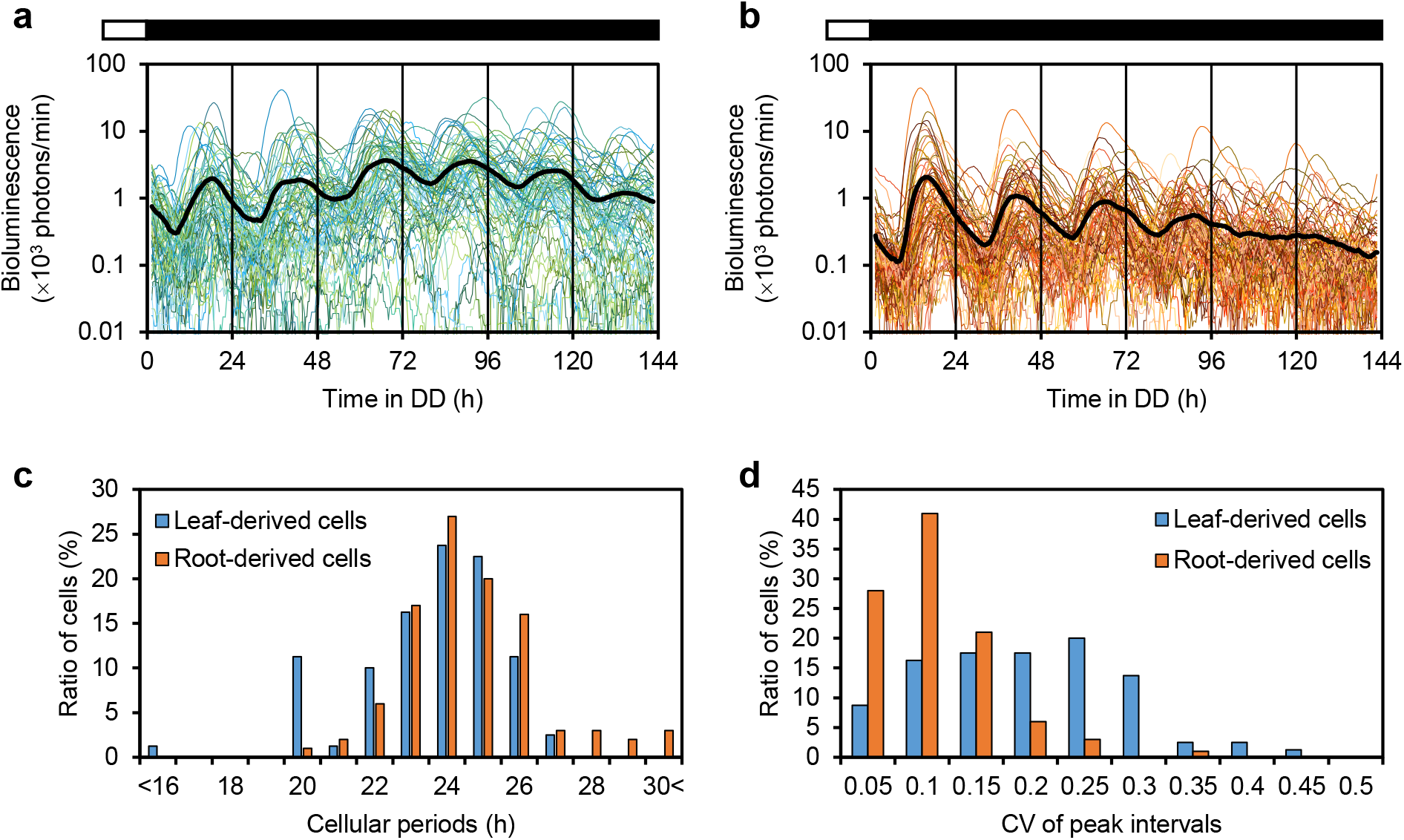
Self-sustained circadian rhythms of leaf- and root-derived cells under standard conditions Protoplasts were isolated from the leaves and roots of *CCA1::LUC* transgenic *Arabidopsis* plants that were entrained to two LD cycles. **a**,**b**, Luminescence traces of individual cells with circadian rhythm [colored lines: 80 leaf-derived cells (**a**); 100 root-derived cells (**b**)] and their mean luminescence (black line) under standard conditions. Open and black boxes indicate light and dark conditions, respectively. **c**, Frequency distribution of cellular periods. **d**, Frequency distribution of the coefficient of variation (CV) in peak intervals.

### Temperature compensation of cellular rhythms within an appropriate temperature range

To confirm the fundamental property of the cellular clock, we determined the temperature compensation of periods. The cells were transferred to DD at 27, 22, 17, or 12°C after protoplast isolation. For both leaf- and root-derived cells, cellular circadian rhythms looked most robust at 17°C and seemed unstable at 12°C (Supplementary Figs. 6, 7). Accordingly, the proportions of cells in the three groups were different at the four different temperatures (Supplementary Fig. 8). The data of peak intervals of every analyzable cell (all in the three groups) at 27, 22, 17, and 12°C are shown in Fig 2. Most peak intervals of bioluminescence rhythms at any temperature were distributed around 24 h during the monitoring (Supplementary Fig. 9). Thus, circadian rhythmicity was maintained in most cells at a single-cell level between 12 and 27°C. At any temperature, the mean values of peak intervals of leaf- and root-derived cells were 23–29 h, while lower temperature resulted in slightly longer period lengths (Fig. 2c,d and Table 2). To evaluate the degree of temperature compensation, we calculated the Q_10_ values. Q_10_ is defined as the ratio of the reaction rates [frequency (1/period) for a rhythm] measured at two temperatures differing by 10°C (Table 2). Q_10_ values of the frequency calculated by every 5°C difference for both leaf- and root-derived cells were 0.98–1.20, indicating that temperature compensation was preserved in the temperature range. Meanwhile, in the range of optimal temperature conditions (17–22°C), the Q_10_ value was larger in the leaf-derived cells than that in the root-derived cells, suggesting that the root-derived cells may have a stricter compensation system for the pace of the endogenous clock than leaf-derived cells. (Table2).

**Table 2.**
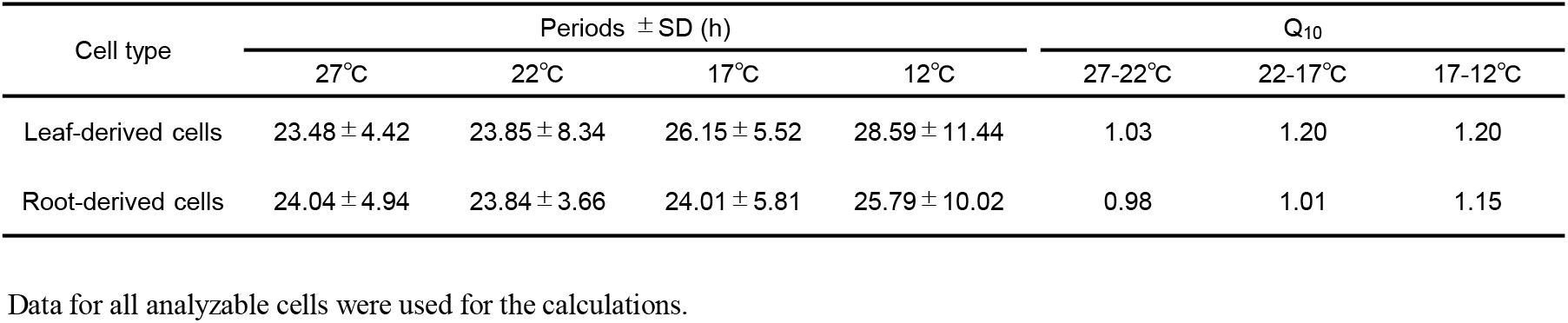
Circadian periods (mean peak intervals) of protoplast-derived cells at 27°C, 22°C, 17°C, and 12°C

| Cell type | Periods ± SD (h) | | | | Q<sub>10</sub> | | |
| --- | --- | --- | --- | --- | --- | --- | --- |
|  | 27°C | 22°C | 17°C | 12°C | 27-22°C | 22-17°C | 17-12°C |
| Leaf-derived cells | 23.48 ± 4.42 | 23.85 ± 8.34 | 26.15 ± 5.52 | 28.59 ± 11.44 | 1.03 | 1.20 | 1.20 |
| Root-derived cells | 24.04 ± 4.94 | 23.84 ± 3.66 | 24.01 ± 5.81 | 25.79 ± 10.02 | 0.98 | 1.01 | 1.15 |

Data for all analyzable cells were used for the calculations.

**Fig. 2.**
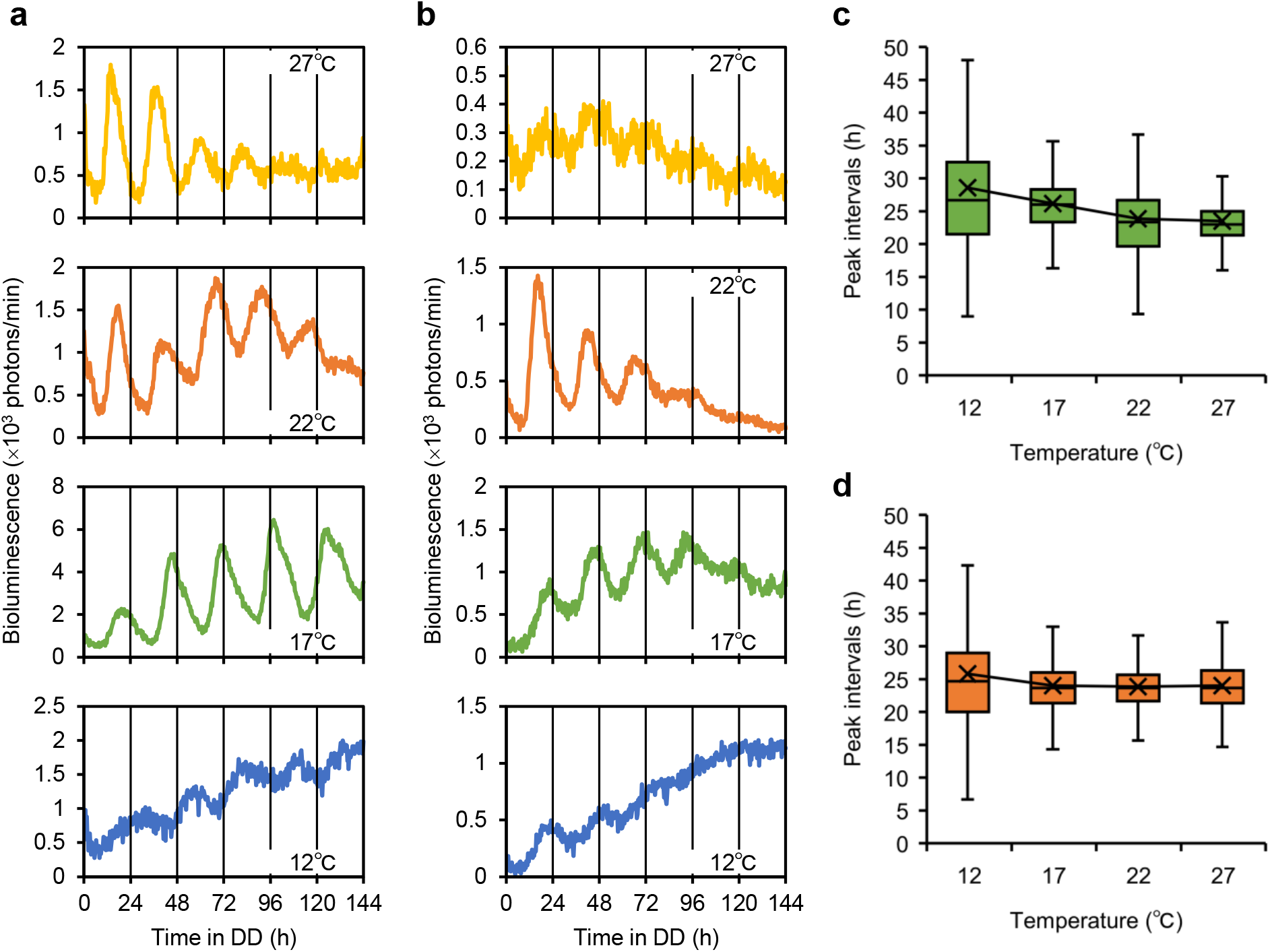
Temperature-compensated periodicity of circadian rhythms of leaf- and root-derived cells Protoplasts were isolated from the leaves and roots of *CCA1::LUC* transgenic *Arabidopsis* plants which were entrained to two LD cycles at 22°C, after which the cells were exposed to 27°C, 22°C, 17°C, or 12°C in constant dark condition for bioluminescence monitoring. **a**,**b**, Mean luminescence (27°C: yellow line, 22°C: orange line, 17°C: green line, and 12°C: blue line) of cellular rhythms of all analyzable cells in leaf-derived cells (**a**) and root-derived cells (**b**). **c**,**d**, Mean peak intervals at each temperature for leaf-derived cells (**c**) and root-derived cells (**d**). In box plots, the center line is the medium; box edges show 25th and 75th percentiles, and whickers extend to minimum and maximum values.

Although a trace of the circadian rhythmicity of the mean luminescence of protoplast-derived cells was observed at 12°C, their amplitudes were quite low or damping (Fig. 2a,b). The peak intervals of individual cellular rhythms showed a wide distribution at 12°C, and hence, the instability of cellular rhythms at this low temperature contributed to the low amplitudes (Fig. 2c,d). Interestingly, the cells that showed skipping or quite low-level peaks between two obvious peaks were observed at 12°C (Supplementary Figs. 6, 7). We compared the synchrony of the initial phases between 12°C and other temperatures (Supplementary Fig. 10). Leaf-derived cells showed a lower synchrony at 12°C, suggesting that low temperature altered the synchronization behavior after protoplast isolation. In contrast, root-derived cells at 12°C showed a similar synchrony of the initial phases to those at higher temperatures. Root-derived cells may have a robust circadian system at low temperatures in terms of precise entrainment. Although the circadian rhythms of root-derived cells showed relatively high synchronization at 12°C, they were promptly desynchronized during monitoring due to the instability of the rhythms. Taken together, protoplast-derived cells maintained circadian periodicity, such that the period length was ∼24 h even when cellular rhythms were highly impaired at low temperature.

### Entrainability of individual cellular rhythms to a light stimulus

As a fundamental property of circadian rhythm, we examined the entrainability of cellular phases to a light stimulus. As illustrated in Supplementary Fig. 11a, the phase transition curves (PTCs) were created by the peak times before and after a light pulse (old phases and new phases). The PTC shows the response of an oscillator to the light stimulus. Because cellular rhythms exhibited a wide variation in the cellular period and initial phase (Fig. 1c and Supplementary Fig. 3), various phases emerged in cells in the monitoring. We exposed the protoplast-derived cells to a 1-hour light pulse at 60 or 72 h after the light-off (Supplementary Fig. 11). For leaf-derived cells treated with a light pulse, the peak times of most of the cellular rhythms appeared 16–30 h after the pulse irrespective of their phases of pulse (Supplementary Fig. 11b). The cellular rhythms showed a Type-0 phase response, which indicated a strong resetting manner by the stimulus^33^. A reference diagram without a light pulse showed a widely scattered plot, and the slope of the linear approximation was near 1 in leaf-derived cells (Supplementary Fig. 11c). This slope was a contrast to that of the PTC for the light pulse. For root-derived cells treated with a light pulse, the peak times of most cellular rhythms appeared 12–27 h after the pulse (Supplementary Fig. 11d). Although the phase after the light pulse appeared to be more advanced in root-derived cells than that in leaf-derived cells, the PTC for both types of cells was similar. The reference diagrams without a light pulse showed widely scattered plots, and the slopes of the linear approximation were near 1 in both leaf- and root-derived cells (Supplementary Fig. 11c,e). These slopes were a contrast to those of the PTCs for the light pulse. Thus, protoplast-derived cells showed an adequate entrainability in response to light stimulus.

Furthermore, we investigated the manner of the entrainment of the cellular rhythms to LD cycles. Almost all the peak times for both leaf- and root-derived cells appeared in the 12-hour light period, suggesting that the cells were entrained to the LD cycles (Fig. 3a,b). The degrees of synchrony for leaf- and root-derived cells were 0.76 and 0.85, respectively (Fig. 3c). The lower degree of synchrony of leaf-derived cells might be due to its less stable periodicity, compared with that of root-derived cells (Fig. 1d). Taken together, the circadian clock with the three fundamental properties was preserved in leaf- and root-derived cells.

**Fig. 3.**
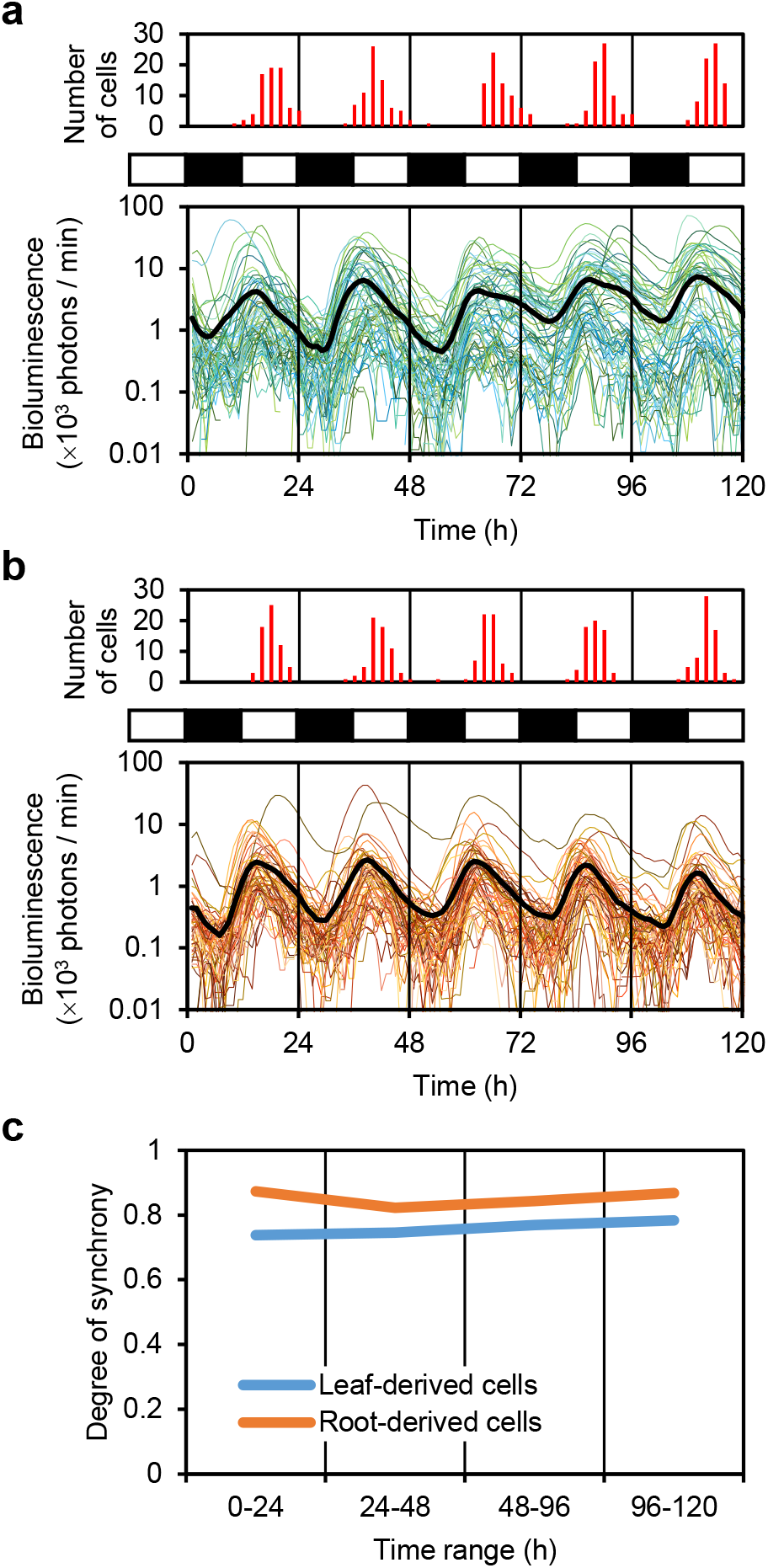
Entrainability of circadian rhythms of protoplast-derived cells to light-dark cycles Protoplasts were isolated from the leaves and roots of *CCA1::LUC* transgenic *Arabidopsis* plants which were entrained to two LD cycles. **a**,**b**, Luminescence traces of individual cells with circadian rhythm (colored lines: 73 leaf-derived cells (**a**); 63 root-derived cells (**b**)) and their mean luminescence (black line) under light-dark conditions. The number of cells peaking at every 2 h is shown at the top. Open and black boxes indicate light and dark conditions, respectively. **c**, Plots for the degrees of synchrony of the peak times of leaf-derived cells (blue line) and root-derived cells (orange line) for each day.

It is interesting to note that all the root-derived cells exposed to LD tended to sustain the bioluminescence rhythm at a similar level during the monitoring (Fig. 3). However, cells exposed to DD tended to show a decrease in the bioluminescence level and a damping rhythm. The circadian clock of the root-derived cells may possibly require light/dark signals for robust oscillation. This is a contrast to the sustainability of circadian rhythms of leaf-derived cells in DD (Fig. 1), suggesting that the circadian oscillation mechanism in leaf-derived cells may be different from that in root-derived cells.

### Increased stability and precision of cellular rhythms of leaf-derived cells in high-cell-density culture

As mentioned earlier, cellular rhythms of leaf-derived cells (cell density in the standard conditions, 2 × 10^4^ cells/4 ml) showed dispersed peak intervals under DD and lower synchrony under LD cycles (Fig. 1d and 3c). Previous studies have shown that the circadian rhythm damped in a diluted culture of protoplasts derived from shoot apex and leaves^12,27^. We assumed that intercellular communication might play a role in the maintenance of coherent rhythms among cells. We first checked the effect of cell density on the circadian rhythms at the cell population level. We co-cultured luminous cells carrying the *CCA1::LUC* reporter with non-luminous wild-type cells at cell density ratios of 1 : 4, 1 : 20, and 1 : 40 of luminous to non-luminous cells (Supplementary Fig. 12). As reported, higher amplitudes of bioluminescence rhythms were sustained with higher cell density under constant light (LL) conditions. In addition, we investigated if the increase in amplitude under high cell density conditions was caused by a mutual synchronization effect on phases of cellular rhythms^34^. We monitored the bioluminescence rhythms of luminous cells (2 × 10^4^ cells/4 ml) that were added to non-luminous cells of the in-phase or anti-phase LD entrainment (4 × 10^5^ cells/4 ml) (Supplementary Fig. 13). Both culture conditions resulted in high-amplitude bioluminescence rhythms with similar phases, suggesting that the effect of high cell density was observed irrespective of the phase information of individual cellular rhythms.

Considering the effect of cell density on amplitude, we assumed that individual cells in a high-cell-density culture might induce a coherent state of circadian oscillations. We approached the mechanism of coherent rhythms by monitoring individual cellular behaviors in the circadian system at a single-cell level. To observe the bioluminescence of single cells dispersedly, we prepared a high-cell-density culture containing luminous cells (2 × 10^4^ cells/4 ml) and non-luminous cells (4 × 10^5^ cells/4 ml). The rhythmicity of leaf-derived cells improved in the high-cell-density culture compared to that in the standard-cell-density culture (Figs. 1 and 4). Supplementary Table 2 describes the proportion of cells in the three categories. The proportion of cells with circadian rhythm (Group 0) was slightly higher in the high-cell-density culture (39%) than in the standard-cell-density culture (32%). The proportion of cells in Group 1 in which rhythmicity disappeared during the monitoring was lower in the high-cell-density culture (11%) than in the standard-cell-density culture (28%) (Supplementary Fig. 8 and Supplementary Table 2). Therefore, high cell density resulted in improved cellular circadian rhythm. Although the bioluminescence rhythm appeared to be more robust in the high-cell-density culture, the cellular periods (high-cell-density: 23.2 ± 1.9 h, standard-cell-density: 23.0 ± 2.2 h) of the leaf-derived cells at the different cell densities were almost the same (Figs. 1c and 4b). Interestingly, the CV of cellular periods was lower (0.13) in the high-cell-density culture than in the standard-cell-density culture (0.17), suggesting that only the stability of each cycle was improved in the high-cell-density culture, without a change in the period length of the cellular rhythm (Figs. 1d and 4c). The difference in the stability of the rhythm under constant conditions was presumed to be induced by a difference in the degrees of internal fluctuations (noise) in the oscillation machinery^8^. For leaf-derived cells, the high-cell-density condition may decrease the degree of internal fluctuations in the cellular circadian clock. A higher synchrony of cellular rhythms under LD cycles was also observed in the high-cell-density culture than in the standard-cell-density culture (Supplementary Fig. 14). A theoretical study revealed that the internal-noise-dependent precision of the phase under LD cycle is a property of self-sustained circadian oscillators with robust protection against external noises from the environment^7^. The circadian clock of leaf-derived cells is possibly this type of self-sustained oscillator.

**Fig. 4.**
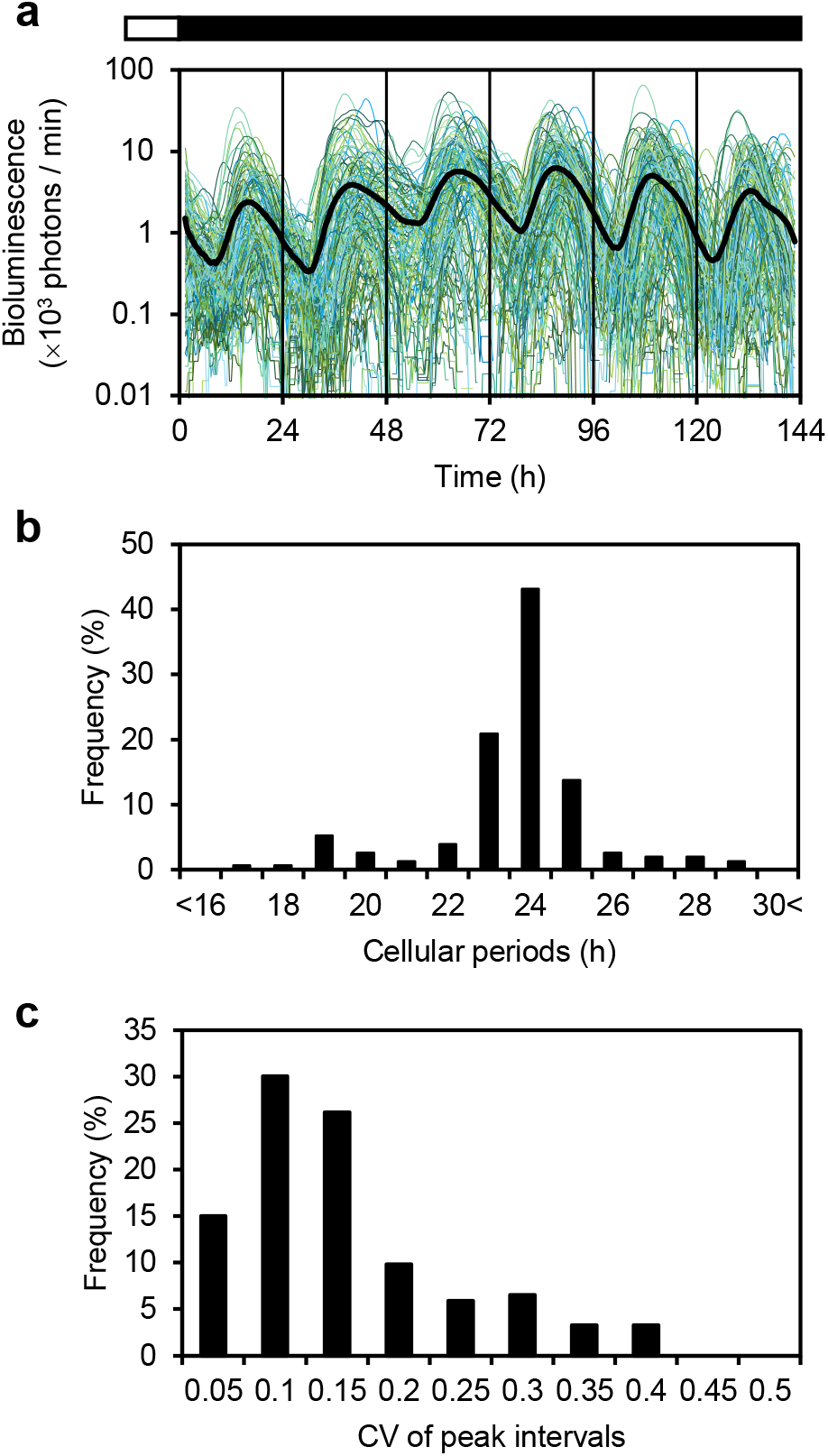
Stabilized cellular rhythms of leaf-derived cells in a high-cell-density culture Protoplasts were isolated from the leaves of *CCA1::LUC* transgenic plants and wild-type plants that were entrained to two light-dark cycles. Luminous cells (2 × 10^4^ cells/4 ml) with non-luminous cells (4 × 10^5^ cells/4 ml) were prepared. **a**, Luminescence traces (colored lines) and mean luminescence (black line) of cells with circadian rhythm (153 leaf-derived cells). Open and black boxes indicate light and dark conditions, respectively. **b**, Frequency distribution of cellular periods. **c**, Frequency distribution of the coefficient of variation (CV) in peak intervals.

The improvement in the stability of each cycle in the high-cell-density culture was dependent on temperature, and it was observed at 27, 22, and 17°C, but not at 12°C (Supplementary Fig. 15). Because cellular rhythms were very unstable at 12°C, the high cell density might have influenced only cells completing each circadian cycle. Interestingly, at each temperature, the mean values of the peak intervals of cellular rhythms were almost the same irrespective of cell densities. This supports the findings that indicate that the stability of each cycle was improved in the high-cell-density culture, without any changes in the period length of the cellular rhythm. Thus, the natural frequency of cellular clocks is possibly not related to the stability of each cycle.

### Increased stability of cellular rhythms of leaf-derived cells in a conditioned medium

The effect of cell density on cellular rhythms is regarded as a social effect in culture medium. To explore the associated mechanism, we investigated the effect of conditioned medium without cells on the stability of cellular rhythms. We used a conditioned medium from a 1-week high-cell-density culture of leaf-derived cells. Leaf-derived cells in the conditioned medium at the standard cell density showed robust circadian rhythms (mean period: 23.3 ± 2.3 h) with a low CV value (0.12), similarly to those in the high-cell-density culture (Fig. 4 and Supplementary Fig. 16). Thus, the effect exerted by high cell density is likely associated with a higher concentration of substances secreted by the culture cells. In addition to the stability of each cycle, the high cell density and the conditioned medium affected the amplitude of cellular rhythms (Supplementary Fig. 17). In the standard-cell-density culture, the amplitude gradually decreased; however, the amplitude slightly increased in the high-cell-density culture and remained constant in the conditioned medium. These results indicated that the social effects were mediated by substances secreted by the cells, which induced coherent rhythms by maintaining cellular amplitude and increasing cycle-to-cycle stability.

## DISCUSSION

Our single cell monitoring of bioluminescence of leaf- and root-derived cells from an *Arabidopsis CCA1::LUC* transgenic plant revealed the characteristics of cellular circadian behavior under constant conditions, without changes in extracellular environments, and cell-autonomous entrainment properties to light-dark cycles. Although the circadian behavior of individual cells exhibited a large variation in the stability of rhythmicity, as a whole, their circadian rhythms showed the three fundamental properties: ∼24-h periodicity, entrainability, and temperature compensation of the period. Under standard culture conditions, intercellular communication may not occur because the cells are not in contact with other cells in the liquid medium. Thus, it has been clearly demonstrated that a circadian rhythm with the fundamental properties is generated through cell-autonomous machinery in plants. Differences were observed between leaf- and root-derived cells with regard to the stability of cellular rhythms (the magnitude of noise on the rhythm).

Circadian rhythms of isolated cells cultured in W5 liquid medium (inorganic salts and 5% FBS) under DD conditions are likely to reflect the native features of cellular clocks without the influence of light signals and nutrient conditions. Despite the artificial conditions, the circadian features of the cells were similar to those observed in corresponding organs. The mean periods of leaf-derived cells (Supplementary Table. 1) were comparable to those of detached leaves under DD conditions^18,22,27^, whereas those of root-derived cells were shorter than those reported for roots under DD conditions^11,21,24,35^. Interestingly, period lengths are highly variable even for same roots, and the involvement of input signals from neighboring cells in the determination of period has been proposed^24^. The root-derived cells in the culture medium may rarely receive environmental input, and they show period lengths as a native feature of their clocks. The damping oscillation of the root-derived cells (Fig. 1), which was observed in this study, was consistent with a previous report on intact or excised roots^11,12^, suggesting that the tendency of damping in the roots may be attributed to individual cellular clocks. In contrast, leaf-derived cells sustained the amplitudes of the rhythms (Fig. 1), and the sustained amplitudes were also observed in detached leaves under DD conditions^18,22^. In plants, low temperature disrupts circadian rhythmicity^36,37^, and we observed highly unstable circadian rhythms of protoplast-derived cells at 12°C (Fig. 2). However, it has been reported that the *CCA1* transcript showed a high amplitude circadian rhythm in *Arabidopsis* seedlings under constant light (LL) at 12°C^37^. The circadian clocks of isolated cells may be more unstable than those of tissues at low temperature. With respect to entrainability, we confirmed that protoplast-derived cells can be reset by light-dark cycles and single light pulse (Fig. 3 and Supplementary Fig. 11). The precision of the phase under light-dark cycles was higher in root-derived cells than in leaf-derived cells at the standard cell density (Fig. 3). The difference in precision may not be associated with the difference in the organs, but with the difference in the stability of circadian rhythm between the cell types as leaf-derived cells at high cell density showed more stable circadian rhythms and consequently improved precision (Supplementary Fig. 14). In plants, a tissue with high cell density may show higher precision in phasing and more stable circadian rhythms compared to a tissue with low cell density. This idea is consistent with the phenomena in which meristematic tissues and vascular tissues showed more robust rhythms compared with other mature tissues^12,13,38^.

The shoots and roots of a plant are under different environmental conditions; the former is subjected to larger degrees of daily changes and more frequent fluctuations in light and temperature than the latter^39^. Therefore, circadian clock properties of these organs are likely to be differently tuned to adapt to the environments of their habitats. In fact, it was reported that the circadian clocks of shoots and roots have different rhythmic properties^11,12,40^. The root clocks respond to light more strongly than the leaf clocks^40^, and the high sensitivity should be more beneficial to the circadian system in the belowground than that in the aboveground. The damped oscillations observed in the root-derived cells may be responsible for the higher sensitivity to light (Figs. 1 and 3). This may be attributed to a decrease in the amplitude of the rhythm, which resulted in a larger phase shift by strength of perturbation from the environment in the limit cycle model for the self-sustained circadian clock^41^. In contrast, high amplitude, which was observed in leaf-derived cells, confers robust protection on oscillation against environmental noises (Fig. 1). Therefore, leaf- and root-derived cells harbor circadian clocks that exhibit adaptive mechanisms to local environments. This suggests that circadian properties specific for an organ in plants may primarily arise from the circadian nature of individual composing cells. It is interesting to note that biochemical clocks face a trade-off between internal and external noise resistances; for example, a circadian oscillation with high amplitude shows the precision of day-night entrainment against environmental noises, and the precision is liable to depend on the internal molecular noise, such as fluctuations of clock components^7^. The imprecise entrainment of leaf-derived cells under standard-cell-density conditions may be caused by a high degree of internal noise as the cells were cultured under constant physical conditions. The improved precision observed with high cell density may be due to a reduction in internal noise. Such differences in the degree of internal noise could be associated with tissue/organ-specific circadian properties in plants.

## MATERIALS AND METHODS

### Plant materials and growth conditions

Wild-type (Col-0) and a transgenic *CCA1::LUC* (Col-0 background) *Arabidopsis thaliana* plants were grown in plates containing 0.5×Murashige and Skoog salts, 0.8% agar and 1% sucrose^42^ at 22°C under LL conditions (30−36 µmol m^-2^ s^-1^). Before sowing, surface-sterilized seeds were stratified for 2 days at 4°C. Cell lines carrying *CCA1::LUC* have been described previously ^28,29^.

### Protoplast isolation and culture

Protoplasts were isolated from the leaves and roots of 3–4-week-old plants as previously described^27,30^ with minor modifications. Before protoplast isolation, the plants were entrained (synchronized) by exposure to two 12 h light/12 h dark (LD) cycles. After 3–4 h of exposure to light, fifty leaves or 15 roots were detached from the plants. Protoplasts were isolated using enzyme solution, and the released protoplasts were filtered through a sterile 70 µm nylon mesh and harvested by centrifugation (leaf-derived cells at 500 g for 2 min; root-derived cells at 1200 g for 5 min) at 22°C. The protoplast pellet was resuspended in W5 solution and more protoplasts were harvested by centrifugation (leaf-derived cells: at 500 g for 3 min; root-derived cells: at 1200 g for 7 min). The resulting protoplast pellet was resuspended and more protoplasts were harvested by centrifugation (leaf-derived cells: at 100 g for 3 min; root-derived cells: at 800 g for 5 min). The isolated protoplasts were cultured in W5 solution supplemented with 5% fetal bovine serum (FBS) (BioSource International, Camarillo, CA; BioWest, Nuaillé, France) and 0.1 mM luciferin, containing D-luciferin and potassium salt (Goryochemical, Japan).

### Single-cell bioluminescence monitoring

Single-cell bioluminescence imaging was performed as previously described^20,43,44^, with modifications, to monitor the protoplast-derived (isolated) cells. A cooled electron-multiplying charge-coupled-device (EM-CCD) camera (ImagEM C9100-13; Hamamatsu Photonics, http://www.hamamatsu.com/) under a macro zoom microscope (MVX-10 with an MVPLAPO 2×C lens; Olympus Optical, https://www.olympus-lifescience.com/) was placed in a lightproof box placed in an incubator (KCLP-1000I-CT; NK system, http://www.nihonika.co.jp/), and the temperature was maintained at 27, 22, 17, or 12°C. The EM-CCD camera was controlled by a computer software (HOKAWO, Hamamatsu photonics). White light from a light-emitting diode (LED) light source (RFB2-20SW; CCS Inc., http://www.ccs-grp.com/) controlled by HOKAWO was guided by optical fibers for illumination of the cell culture (7−8 µmol m^-2^ s^-1^ for the 12 h light period; 30−32 µmol m^-2^ s^-1^ for a light pulse). For time-lapse imaging, bioluminescence images were captured with an EM-CCD camera (cooled at −80°C) every 20 and 60 min under DD and LD conditions, respectively. Exposure times for the bioluminescence of leaf- or root-derived cells were for 4 or 6 min, respectively. To remove cosmic ray spikes, two sequential images were captured at a time and the minimum value for each pixel was used for analysis. In the LD cycles, bioluminescence images were captured after 4 min dark treatment for autofluorescence decay. ROIs were manually defined for individual isolated luminous cells, and the bioluminescence intensity in each ROI was quantitated. The luminescence intensity of each ROI was quantified by using the following equation^43^: Number of photons = (signal intensity − background intensity) × 5.8/(1200 × 0.9). Time-series data of the intensities of each ROI were subjected to peak detection. Image analysis and postprocessing were carried out with ImageJ (http://rsbweb.nih.gov/ij/).

### Time series analysis

Peak detection and Fast Fourier transform-nonlinear least squares (FFT-NLLS) were performed using R 4.0.3. (http://r-project.org/) as previously described^20^. The peaks were estimated by a local quadratic curve fitting for the smoothed time series with a 2-h moving average^20^. Period was estimated by two methods: peak intervals and FFT-NLLS analysis. Peak interval was calculated for sequential pairs of peaks. The amplitude of cellular circadian rhythms was obtained by calculating the mean and standard deviation of time-series data from sliding 24-h window. In FFT-NLLS, the rhythm significance is estimated by an RAE that increases from 0 to 1 as the rhythm nears statistical insignificance.

### Bioluminescence monitoring at a cell-population level

Bioluminescence monitoring of total cells in the culture medium in a 35-mm dish was performed as previously described^27,45^. The bioluminescence dish-monitoring system used photomultiplier tubes for bioluminescence detection, and each dish was subjected to measurement for 30 s every 20 min at 22°C. The light intensity of the illumination to the dishes was approximately 30 µmol m^-2^ s^-1^.

### Determination of the degree of synchrony

The degree of synchrony of circadian rhythms in a population (group) of cells for each day was calculated as follows. (1) Plotting the peak time(s) of each cellular rhythm in the 24-h time range on a unit circle and taking each plot as a vector with a magnitude of 1. (2) Calculating the vector sum of the plots. (3) Dividing its magnitude by the number of peaks in the range. The degree of synchrony represents the coherence of the group, ranging from 0 (desynchronized) to 1 (synchronized).

### Preparation of the conditioned medium

Conditioned medium was prepared from a 1-week high-cell-density culture (4 × 10^5^ cells /4 ml) of leaf-derived cells that had been subjected to bioluminescence monitoring under DD conditions. The culture medium was centrifuged at 1000 g for 3 min at 22°C, and the supernatant was filtered through a sterile 0.2 µm cellulose filter. The resulting filtrate was used as the conditioned medium.

### Creation of graphs and plots

Graphs and plots were created with Excel (Microsoft Corporation).

## ACKNOWLEDGEMENTS

We thank Dr. Norihito Nakamichi for experimental materials. We also appreciate Dr. Shogo Ito and Dr. Masaaki Okada for experimental support. This work was supported in part by the Japan Society for the Promotion of Science KAKENHI [Grant numbers JP25650098, JP17KT0022, and JP19H03245 to T.O. and JP19J23441 to S.N.].

---

**Supplementary Fig. 1.**
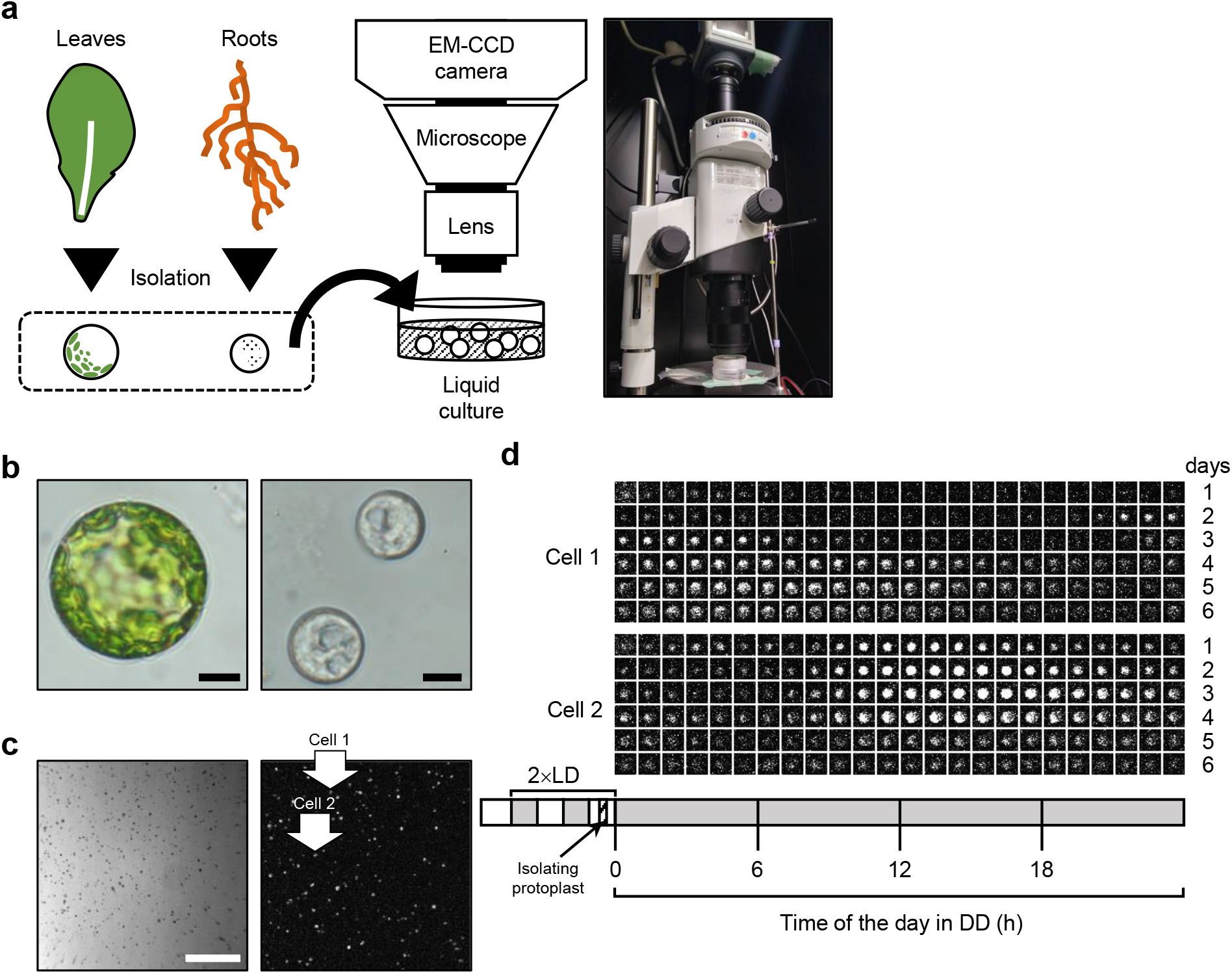
Imaging system for bioluminescence of individual protoplast-derived cells **a**, Schematic overview of the imaging system. Cells were isolated from the leaves and roots as protoplasts and transferred to a culture medium containing 0.1 mM luciferin in a 35-mm dish. Bioluminescence was captured using an Electron Multiplying Charge-Coupled Device camera under a macro zoom microscope. **b**, Protoplasts isolated from leaves (left) and roots (right). Black bars represent 10 μm. **c**, A bright-field image (left) and a luminescence image (right). Bioluminescence of individual cells isolated from the leaves was captured with an exposure time of 240 s at 72 h under a constant dark (DD) condition. A white bar represents 1 cm. **d**, Examples of bioluminescence changes in cells with circadian rhythm (Cell 1 and 2 in panel **c**). The experimental scheme for bioluminescence monitoring is shown in the diagram below. Gray and open boxes indicate dark and light, respectively.

**Supplementary Fig. 2.**
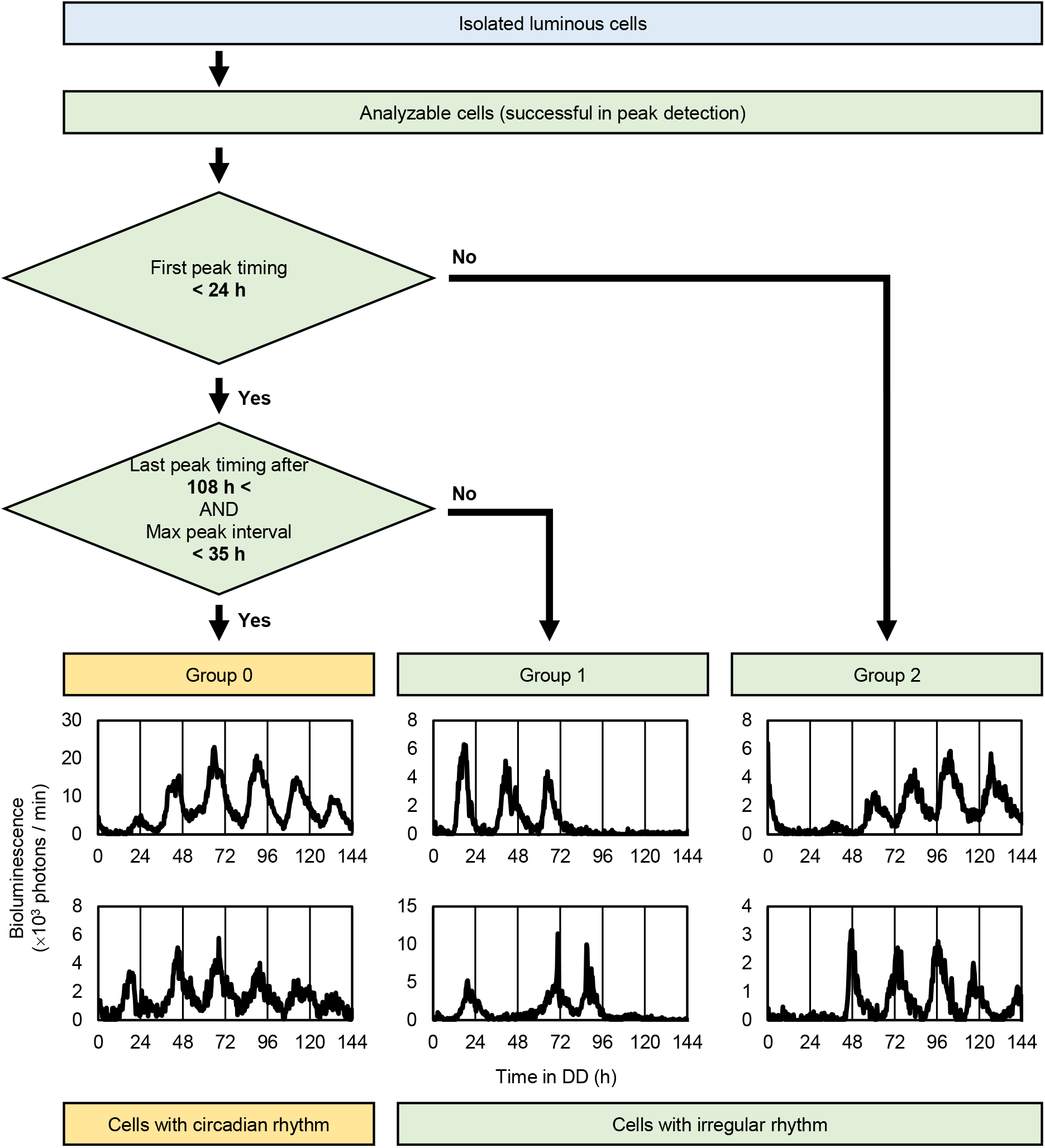
Classification of cells by continuity of bioluminescence rhythms Luminous cells, in isolation from other luminous cells, showing bioluminescence rhythms with detectable peaks were selected for analysis. Cells showing bioluminescence rhythms without a peak in the first 24 h were categorized into Group 2. Of the cells showing bioluminescence rhythms with the first peak in the first 24 h, those with the last peak in a 108−144 h range and with the maximum peak interval at less than 35 h were categorized into Group 0 (cells with circadian rhythm). The rest were categorized into Group 1. Cells in Group 1 and Group 2 had irregular rhythm. Bioluminescence rhythms of cells in each category are presented.

**Supplementary Fig. 3.**
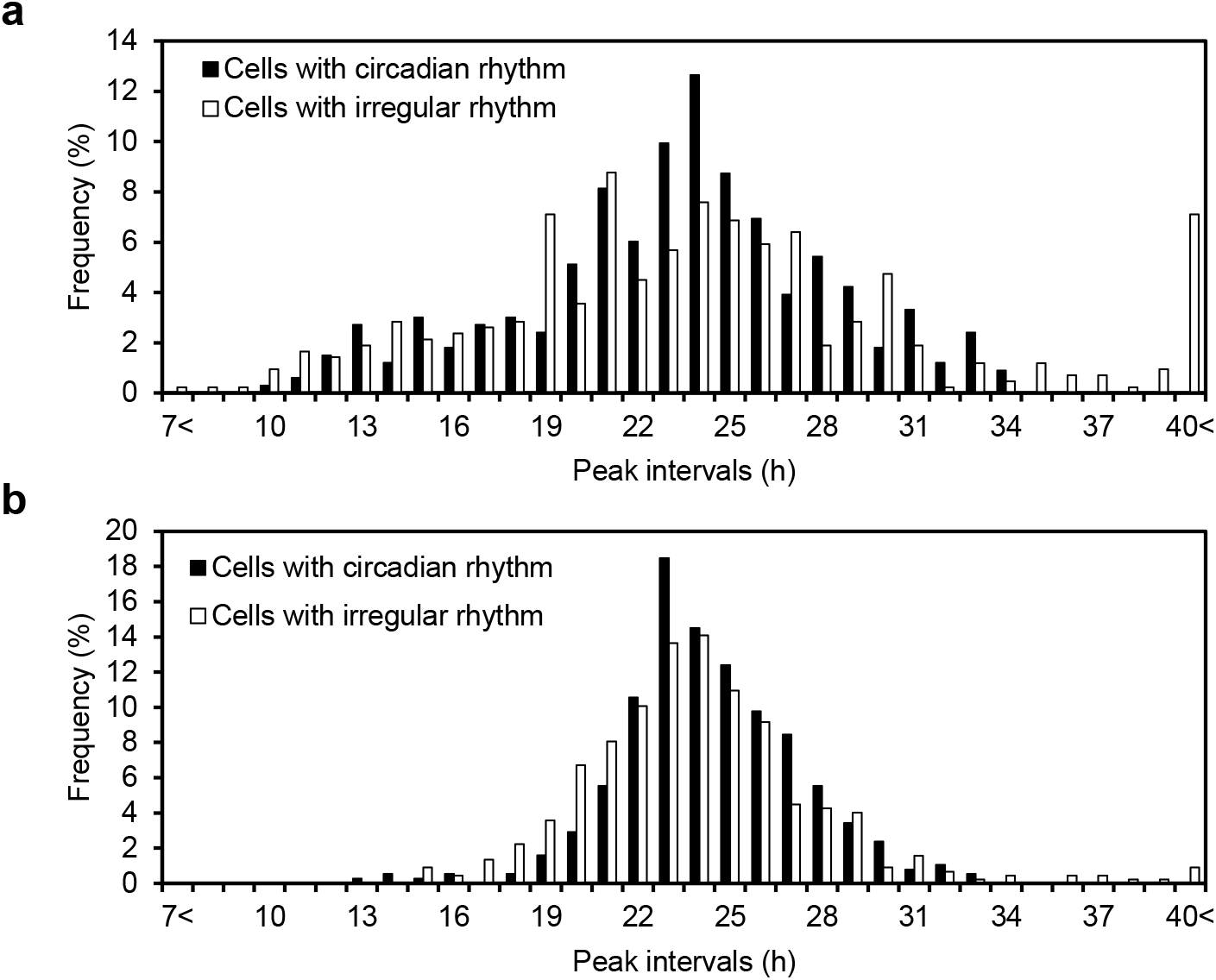
Frequency distributions of peak intervals of bioluminescence rhythms of cells with circadian rhythm and irregular rhythm under standard conditions **a**, Leaf-derived cells. **b**, Root-derived cells.

**Supplementary Fig. 4.**
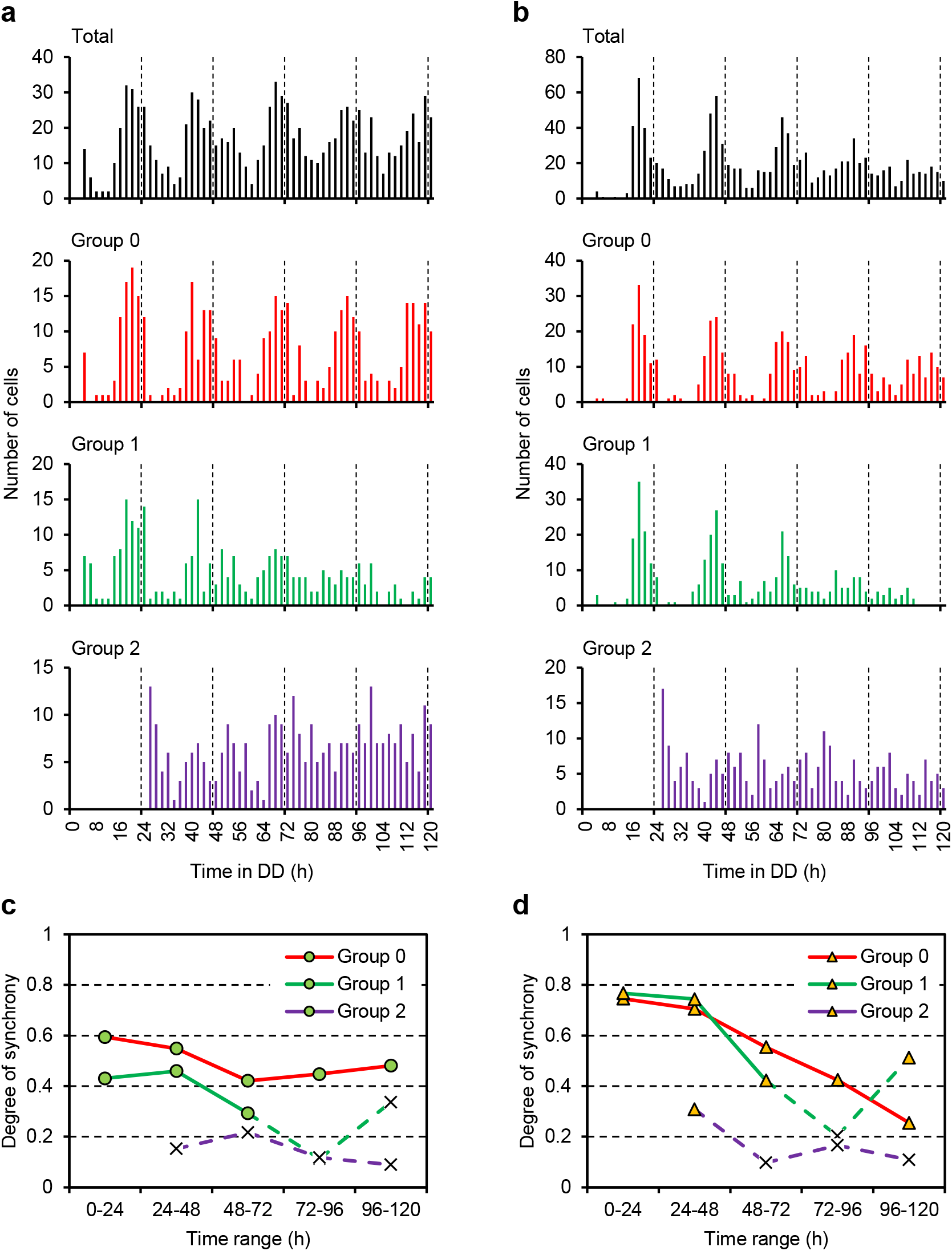
Changes in the degrees of synchrony of cellular rhythms under standard conditions **a**,**b**, Each graph shows the number of cells peaking at every 2 h. Peak times for every cells in Group 0, Group 1, Group 2, for all leaf-derived (**a**) and root-derived (**b**) cells. **c**,**d**, Plots for the degrees of synchrony of peak times in each group (Group 0: red line, Group 1: green line, Group 2: purple line) of leaf-derived (**c**) and root-derived (**d**) cells for each day. Green circle and orange triangle show the statistical significance (Rayleigh test; *p* < 0.01), indicating that the distributions are significantly different from uniform. Cross marks represent non-significant points.

**Supplementary Fig. 5.**
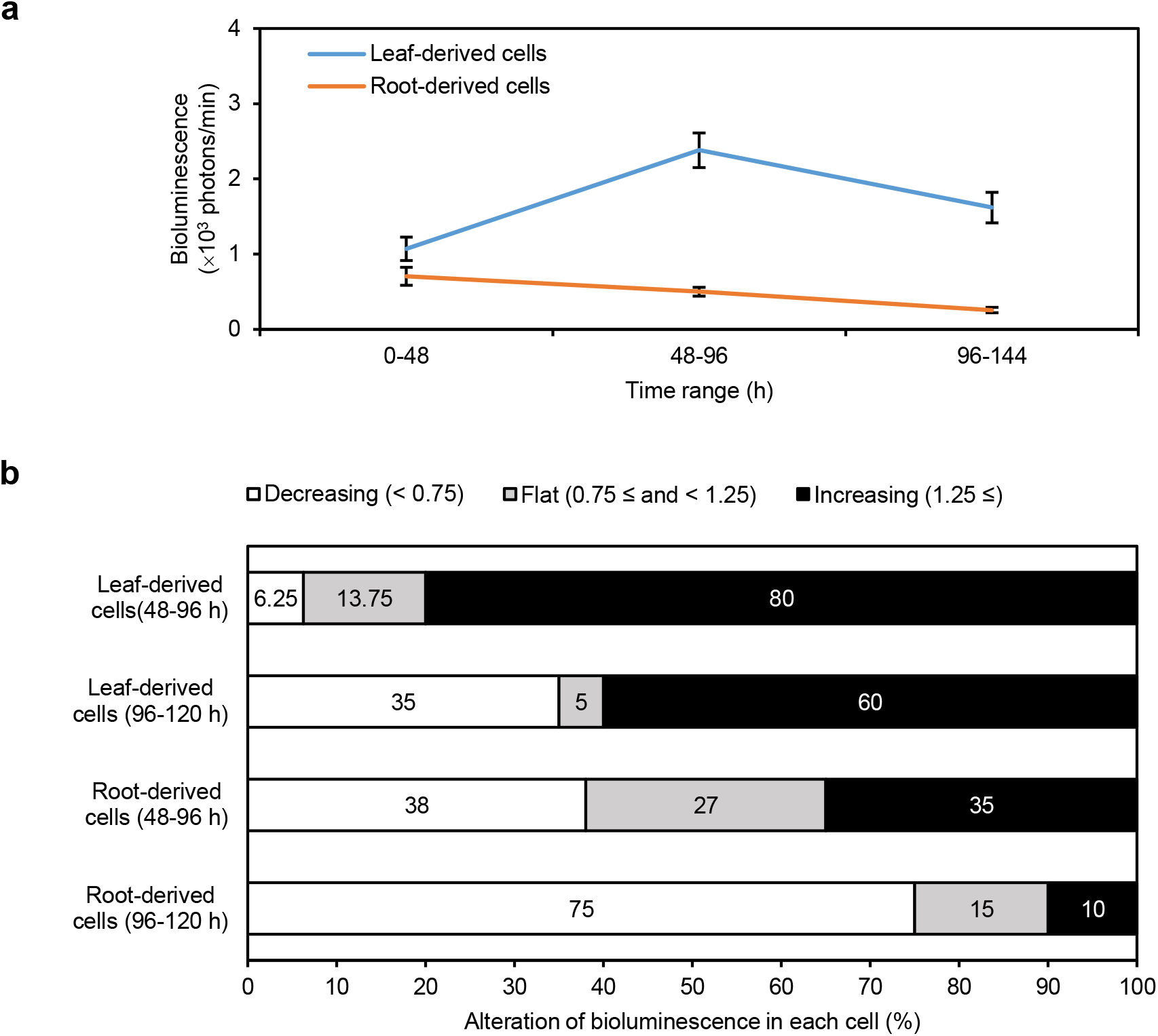
Bioluminescence intensity of leaf- and root-derived cells during monitoring under standard conditions **a**, Mean bioluminescence intensities of leaf-derived (blue line) and root-derived (orange line) cells for 48 h time range. Data are presented as means ± standard error. **b**, 100% stacked bar charts of cells showing decreasing (< 0.75), flat (0.75 ≤ and < 1.25), and increasing (1.25 ≤) bioluminescence between 0–48 h and 48–96 h or between 0–48 h and 96–144 h. Analyzed data are shown in Fig. 1.

**Supplementary Fig. 6.**
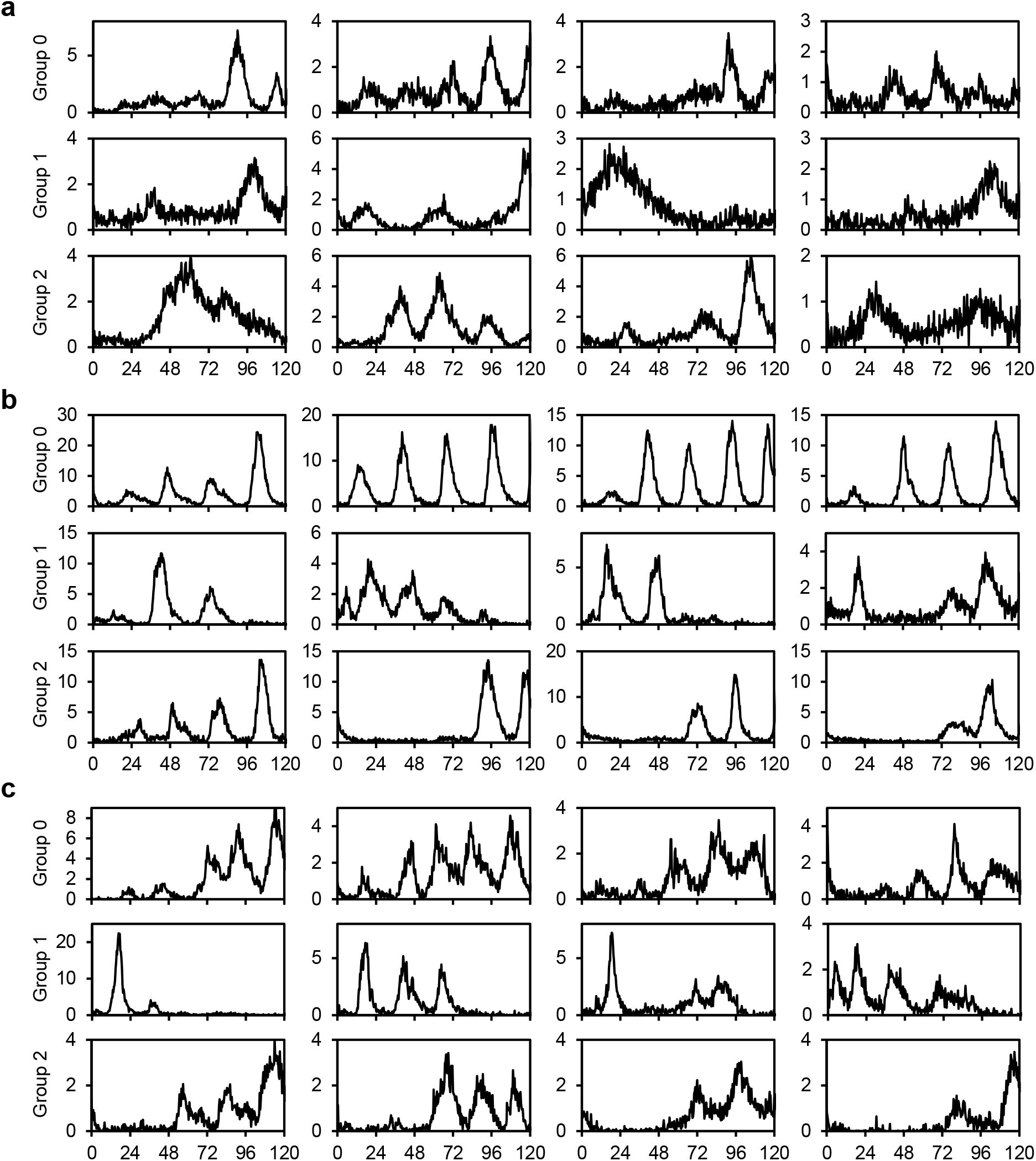
Examples of cellular bioluminescence traces of leaf-derived cells at 12°C, 17°C, and 22°C Four examples for Group 0, Group 1, and Group 2 at 12°C (**a**), 17°C (**b**), and 22°C (**c**)

**Supplementary Fig. 7.**
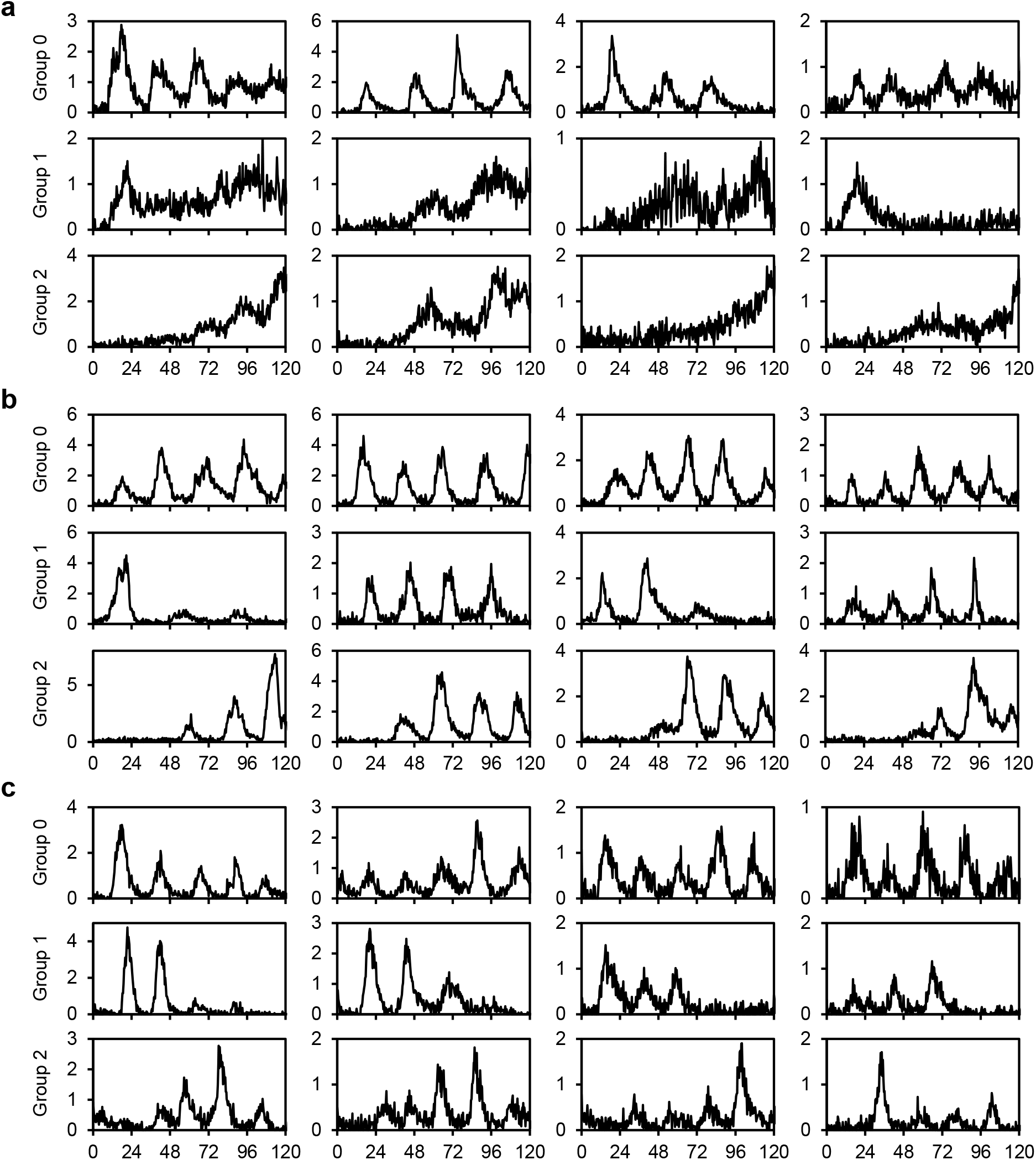
Examples of cellular bioluminescence traces of root-derived cells at 12°C, 17°C, and 22°C Four examples for Group 0, Group 1, and Group 2 at 12°C (**a**), 17°C (**b**), and 22°C (**c**)

**Supplementary Fig. 8.**
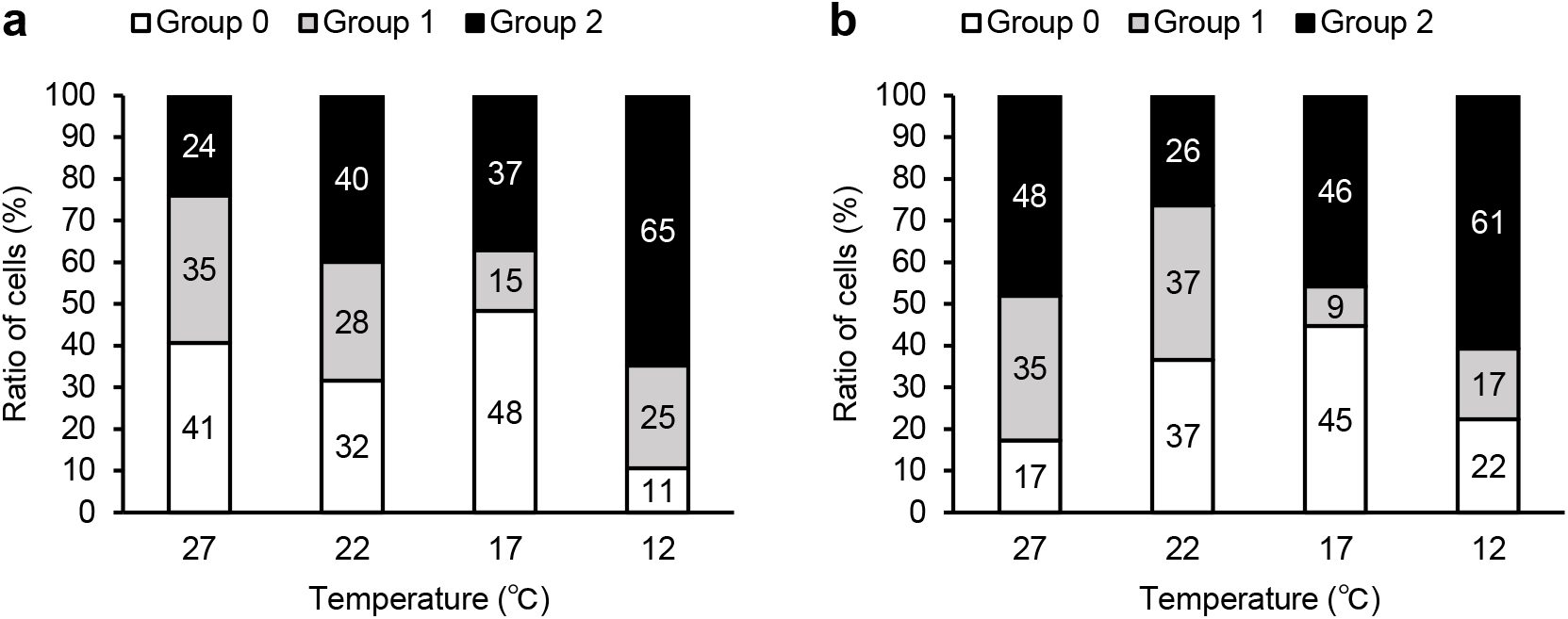
Comparison of the ratios of cells in Group 0, Group 1, and Group 2 at different temperature conditions 100% stacked bar charts of the ratios of cells in the three groups for leaf-derived (**a**) and root-derived (**b**) cells at 27°C, 22°C, 17°C, and 12°C.

**Supplementary Fig. 9.**
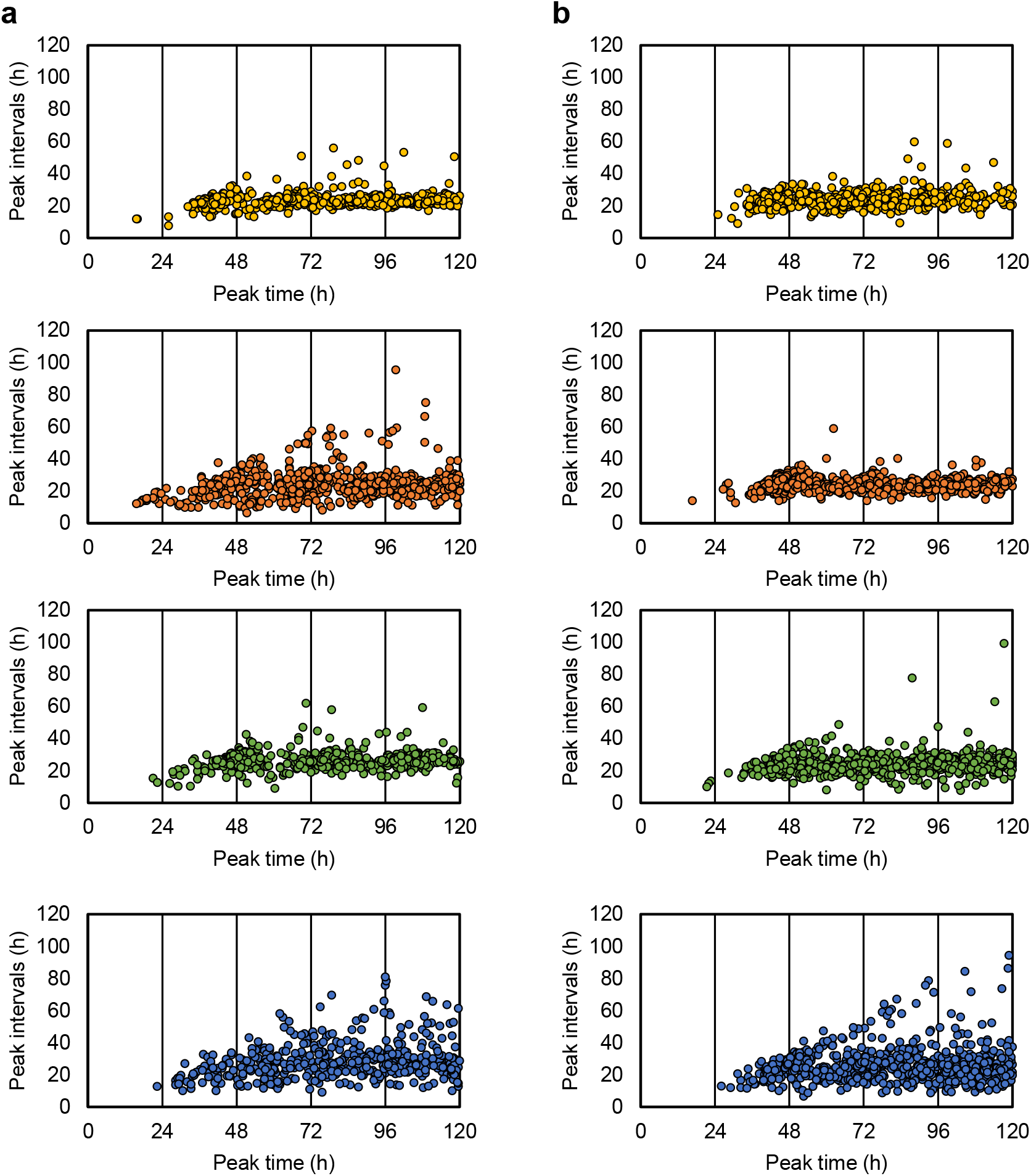
Plots of peak intervals against peak times in the culture of protoplast-derived cells at different temperatures The peak interval at *t*_*k*_ is defined as *t*_*k*_ -*t*_*k-1*_, where time at the *k*th peak of a cellular rhythm is *t*_*k*_ . Plots of peak intervals and peak times of cellular rhythms of all analyzable cells for leaf-derived (**a**) and root-derived (**b**) cells at 27°C (yellow), 22°C (orange), 17°C (green), and 12°C (blue).

**Supplementary Fig. 10.**
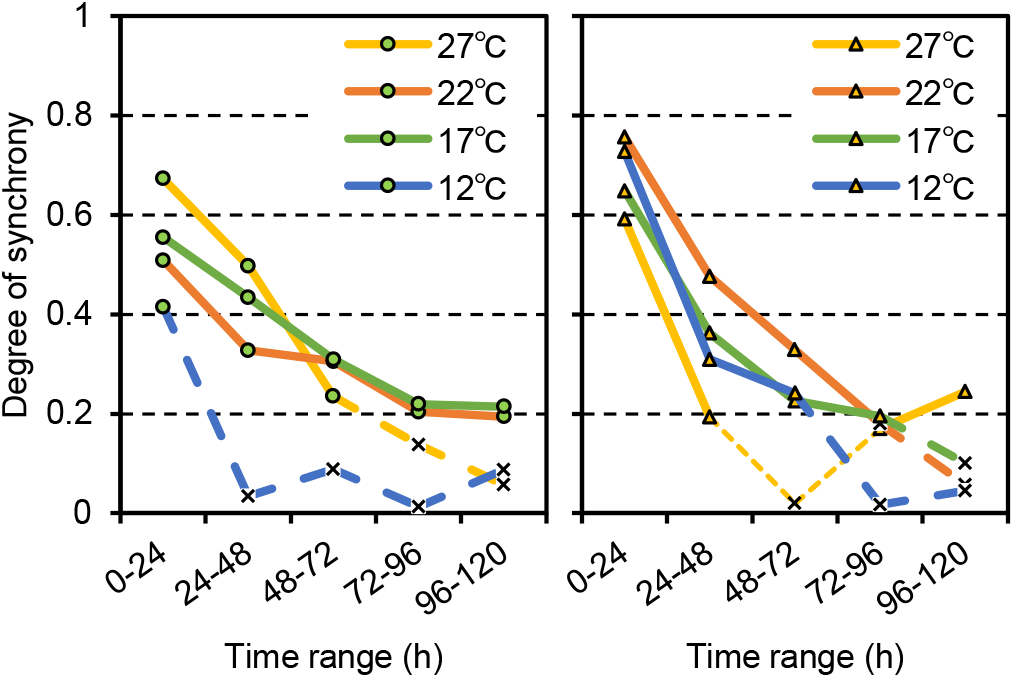
Alteration of the degrees of synchrony of cellular rhythms at different temperatures Plots for the degrees of synchrony of peak times under four temperature conditions (27°C: yellow line, 22°C: orange line, 17°C: green line, and 12°C: blue line) of leaf-derived (left panel) and root-derived (right panel) cells for each day. Green circle and orange triangle show the statistical significance (Rayleigh test; *p* < 0.01), indicating that the distributions were significantly different from uniform. Cross marks represent non-significant points.

**Supplementary Fig. 11.**
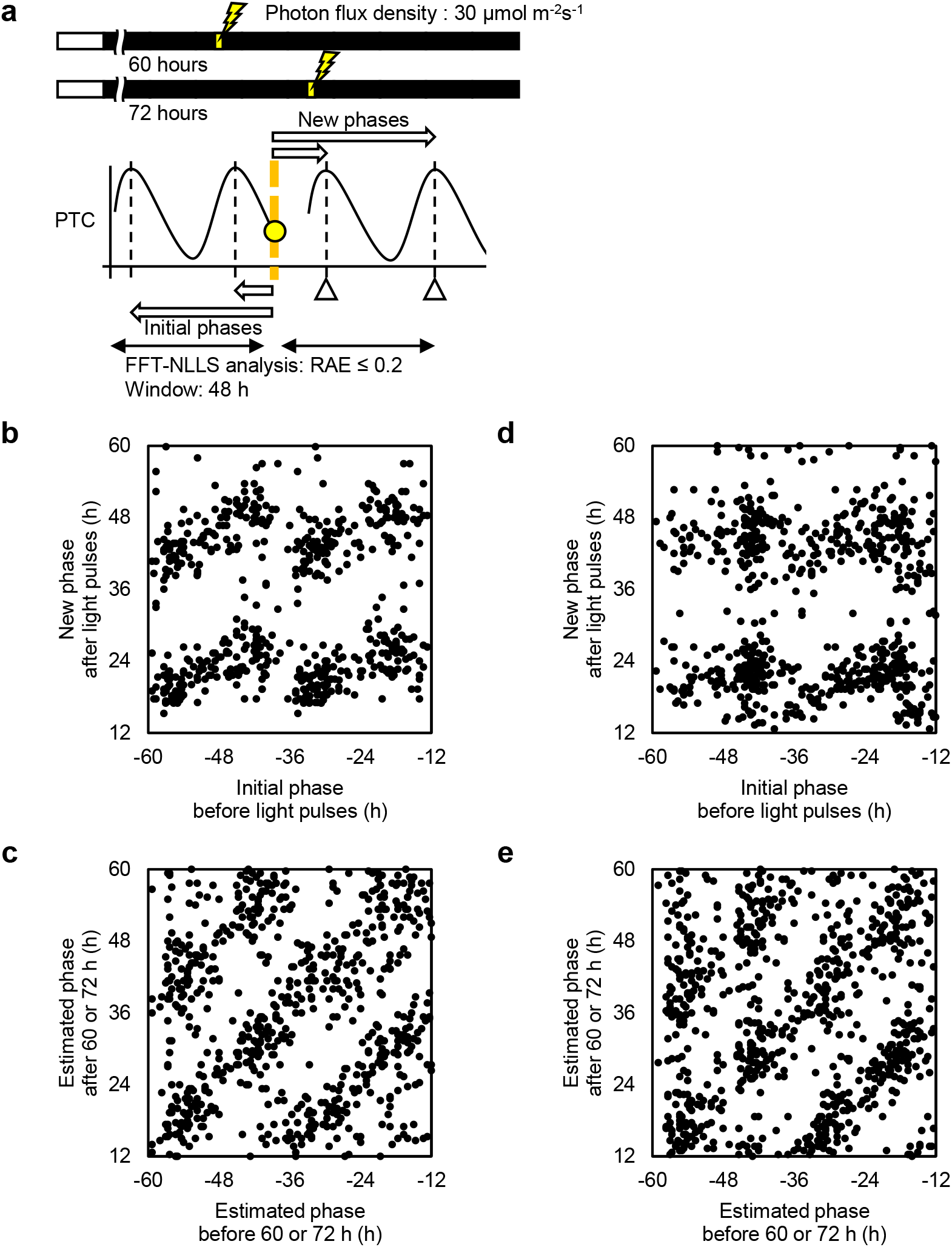
Phase shifts of cellular circadian rhythms to a light pulse **a**, Schematic representation of the experimental strategy to generate a phase transition curve (PTC). An initial phase is a phase of a rhythm at which time the light pulse is given, and in this study, the phase was defined as the subtraction of the time of light pulse from a peak time before the light pulse. A new phase is a phase of the phase-shifted rhythm at the light pulse, and herein, the phase was defined as the subtraction of a peak time after light pulse from the time of light pulse. A PTC represents a plot of initial and new phases for circadian rhythms with various phases. Bioluminescence traces with circadian rhythmicity (RAE ≤ 0.2 in the 48 h ranges before and after the light pulse) were analyzed. A light pulse was given at 60 h or 72 h under constant dark condition. **b**,**c**, PTC for leaf-derived cells (**b**) and a plot representing a reference condition without the light pulse (**c**). **d**,**e**, PTC for root-derived cells (**d**) and a plot representing a reference condition without the light pulse (**e**).

**Supplementary Fig. 12.**
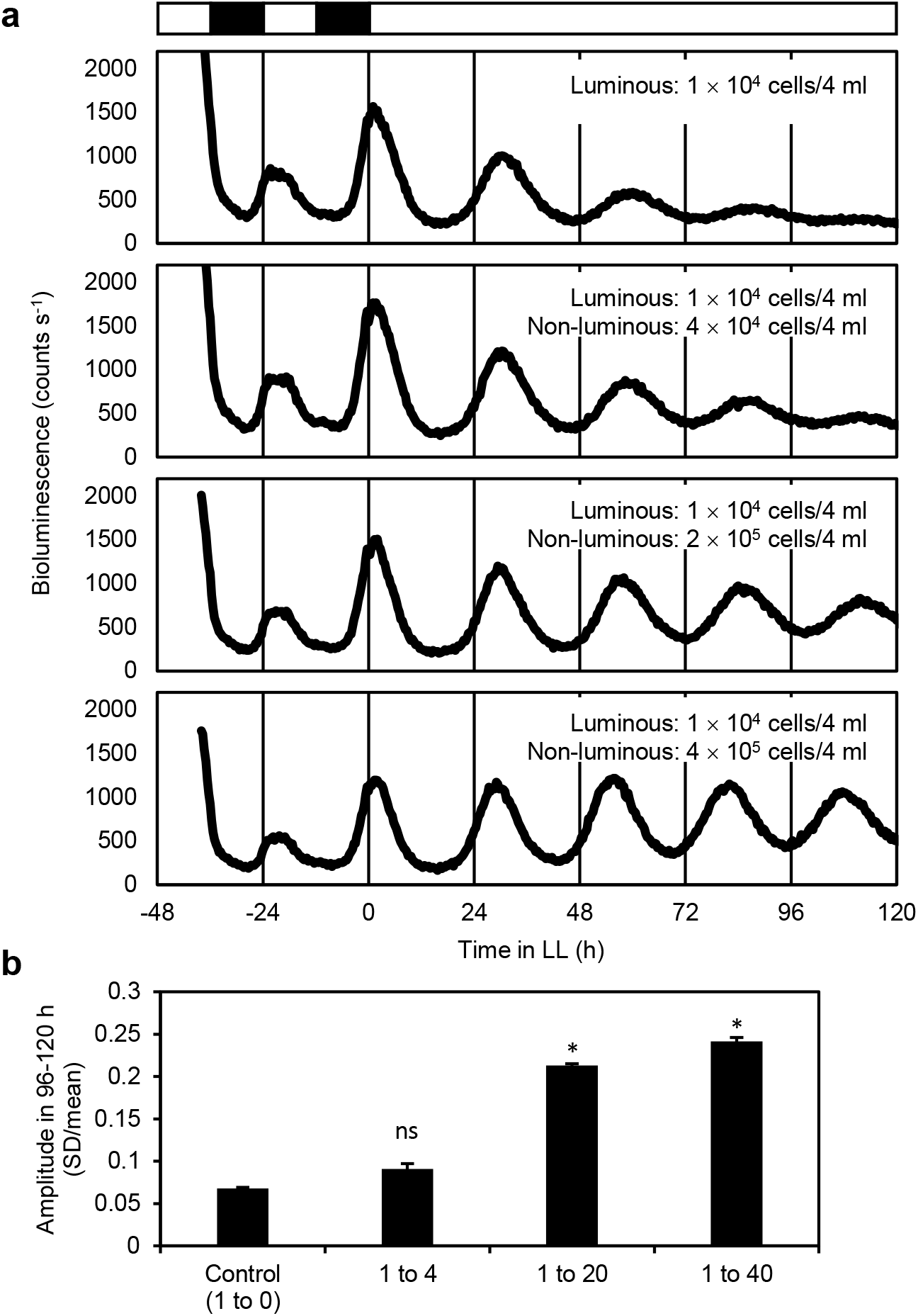
Increasing the cell density in the culture medium increased the amplitude of circadian rhythms of leaf-derived cells at the cell-population level **a**, Luminescence from a culture dish containing leaf-derived cells under constant light conditions at the indicated cell densities of luminous and non-luminous cells. Luminous and non-luminous cells were protoplasts isolated from the leaves of *CCA1::LUC* transgenic plants and wild type plants, respectively. Open and black boxes indicate light and dark conditions, respectively. **b**, Amplitudes of circadian rhythms at each cell density. Amplitude was defined as the coefficient of variation (standard deviation/mean) of bioluminescence intensities at 72 time points in 96−120 h. Data are presented as mean ± standard error for three independent experiments. *, significantly different from the control (*p* < 0.01, using Student’s *t* test). ns = not significant.

**Supplementary Fig. 13.**
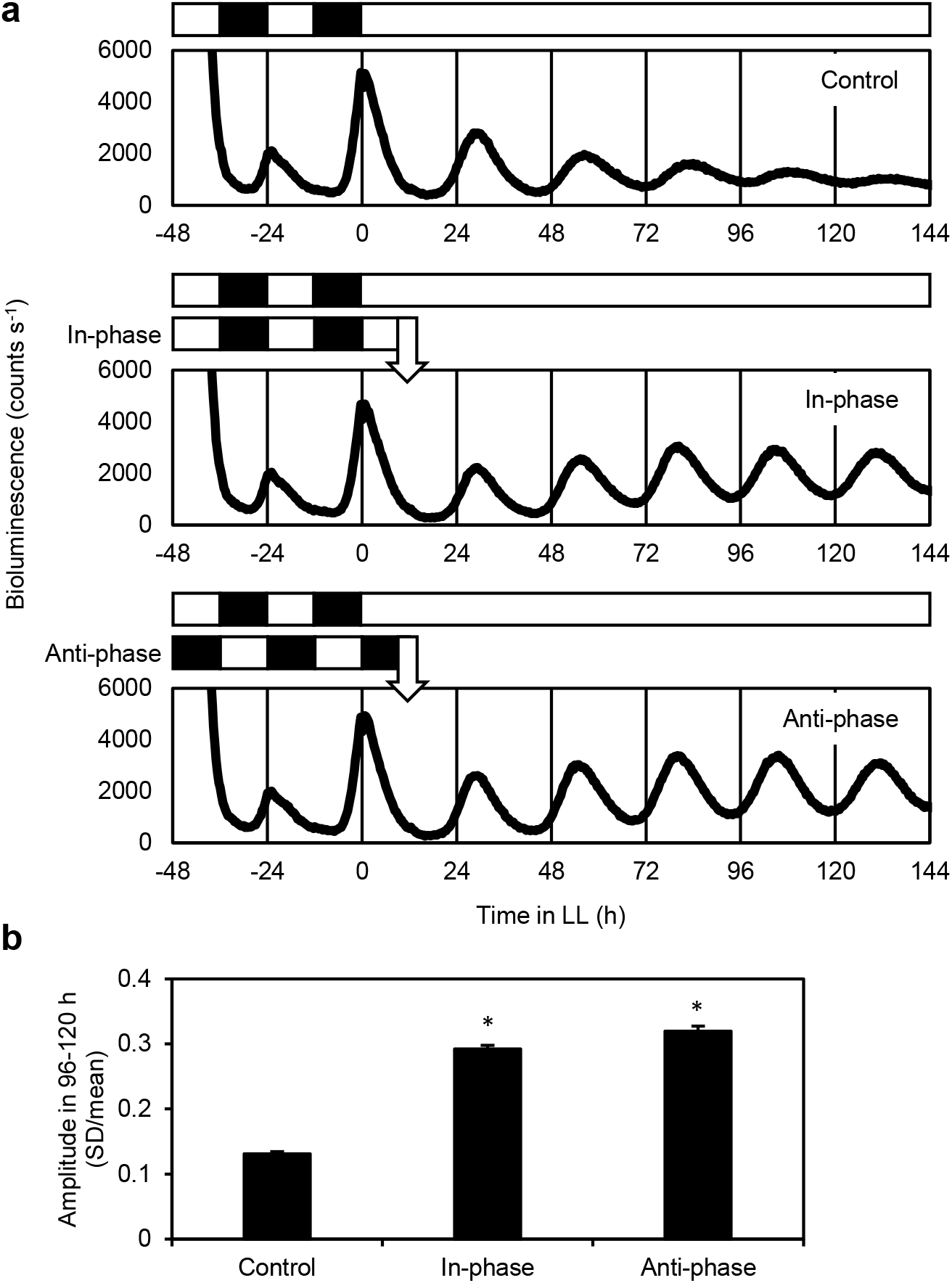
Effects of adding in-phase and anti-phase non-luminous cells on bioluminescence circadian rhythms of leaf-derived cells **a**, Luminescence from a culture dish containing leaf-derived cells under constant light conditions at the cell densities of luminous cells (2 × 10^4^ cells/4 ml) and non-luminous cells (4 × 10^5^ cells/4 ml) . Luminous and non-luminous cells were protoplasts isolated from the leaves of *CCA1::LUC* transgenic plants and wild type plants, respectively. Non-luminous cells were added to the culture dish at the time indicated by an arrow. Open and black boxes indicate light and dark conditions, respectively. **b**, Amplitudes of circadian rhythms. Amplitude was defined as the coefficient of variation (standard deviation/mean) of bioluminescence intensities at 72 time points in 96−120 h. Data are represented as mean ± standard error for three independent experiments. *, significantly different from the control (*p* < 0.01, using Student’s *t* test). The difference in the amplitude between in-phase and anti-phase was not significant (*p* = 0.24).

**Supplementary Fig. 14.**
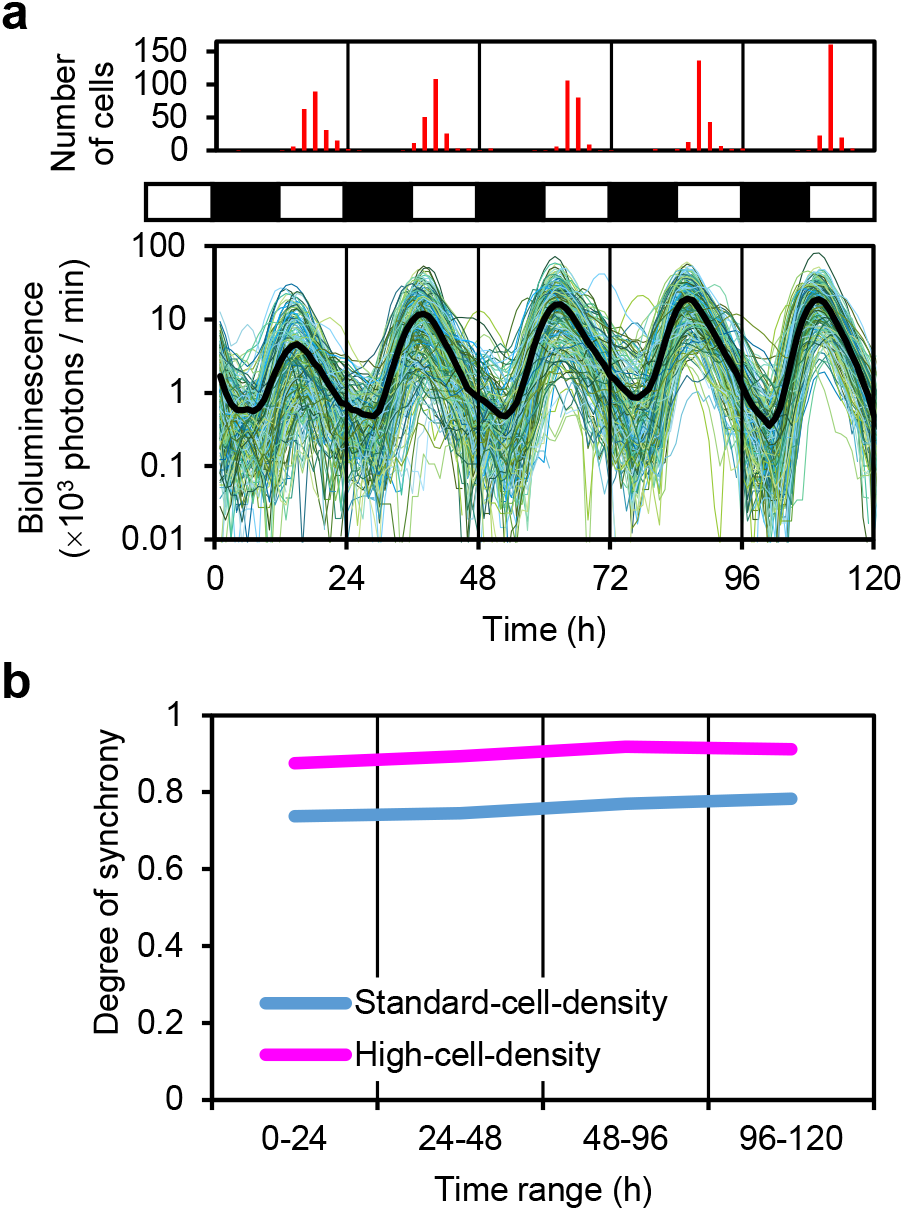
Increased synchrony of leaf-derived cells in the high-cell-density culture under light-dark cycles **a**, Luminous traces of individual cells with circadian rhythm (colored lines: 203 leaf-derived cells) and their mean luminescence (black line) under light-dark cycles in the high-cell-density culture. The number of cells peaking at every 2 h is shown at the top. Open and black boxes indicate light and dark conditions, respectively. **b**, Plots for the degrees of synchrony of peak times of leaf-derived cells in the standard-cell-density culture (blue line) and in the high-cell-density culture (magenta line) for each day.

**Supplementary Fig. 15.**
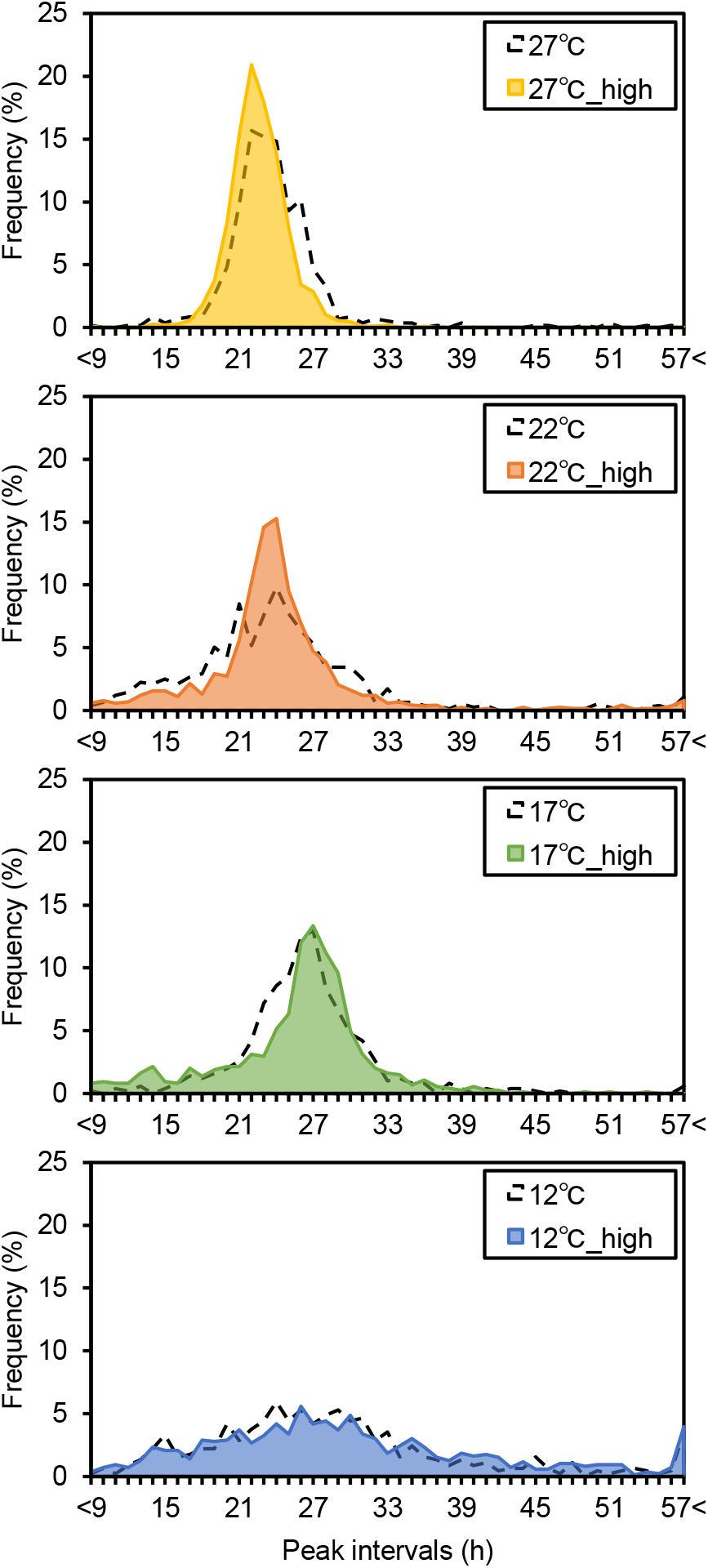
Frequency distributions of peak intervals of leaf-derived cells in high-cell-density culture at different temperatures Protoplasts were isolated from the leaves of *CCA1::LUC* transgenic and wild-type plants and were entrained to two LD cycles, after which the cells were exposed to different temperatures (27°C, 22°C, 17°C, or 12°C) in constant dark for bioluminescence monitoring. Luminous cells (2 × 10^4^ cells/4 ml) and non-luminous cells (4 × 10^5^ cells/4 ml) were prepared. Frequency polygons of peak intervals at each temperature for leaf-derived cells in the high-cell-density culture (27°C: yellow, 22°C: orange, 17°C: green, and 12°C: blue). The broken line in each plot represents the frequency distribution of cells in the standard-cell-density culture (refer Fig. 2c).

**Supplementary Fig. 16.**
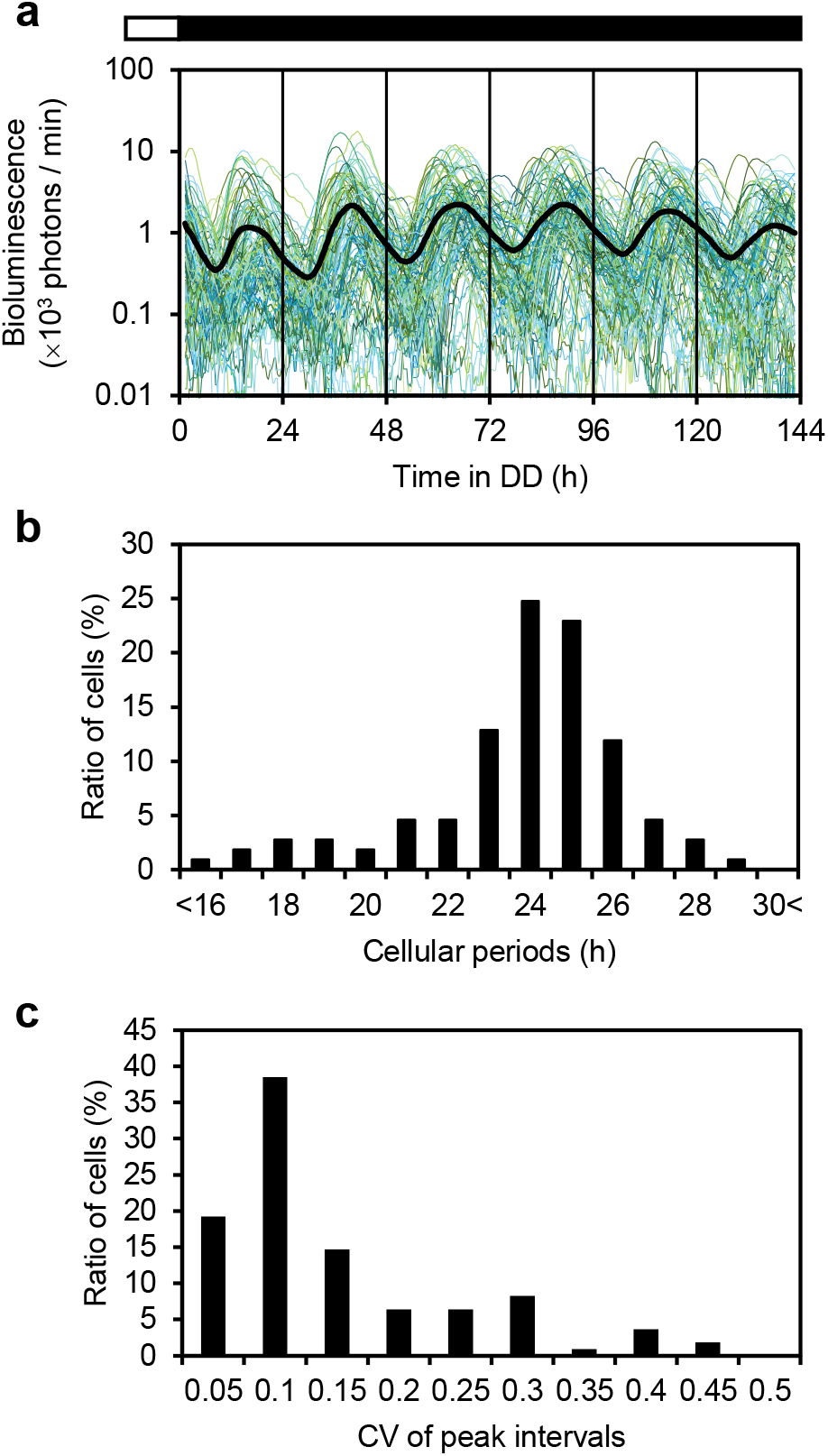
Stabilized cellular rhythms of leaf-derived cells under standard-cell-density conditions with a conditioned medium Protoplasts were isolated from the leaves of *CCA1::LUC* transgenic plants and wild-type plants which were entrained to two LD cycles. Luminous cells (2 × 10^4^ cells/4 ml) were prepared and transferred into a conditioned medium that was prepared from a 1-week high-cell-density culture of leaf-derived cells. **a**, Luminescence traces (colored lines) and mean luminescence (black line) of cells with circadian rhythm (109 leaf-derived cells). Open and black boxes indicate light and dark conditions, respectively. **b**, Frequency distribution of cellular periods. **c**, Frequency distribution of the coefficient of variation (CV) in peak intervals.

**Supplementary Fig. 17.**
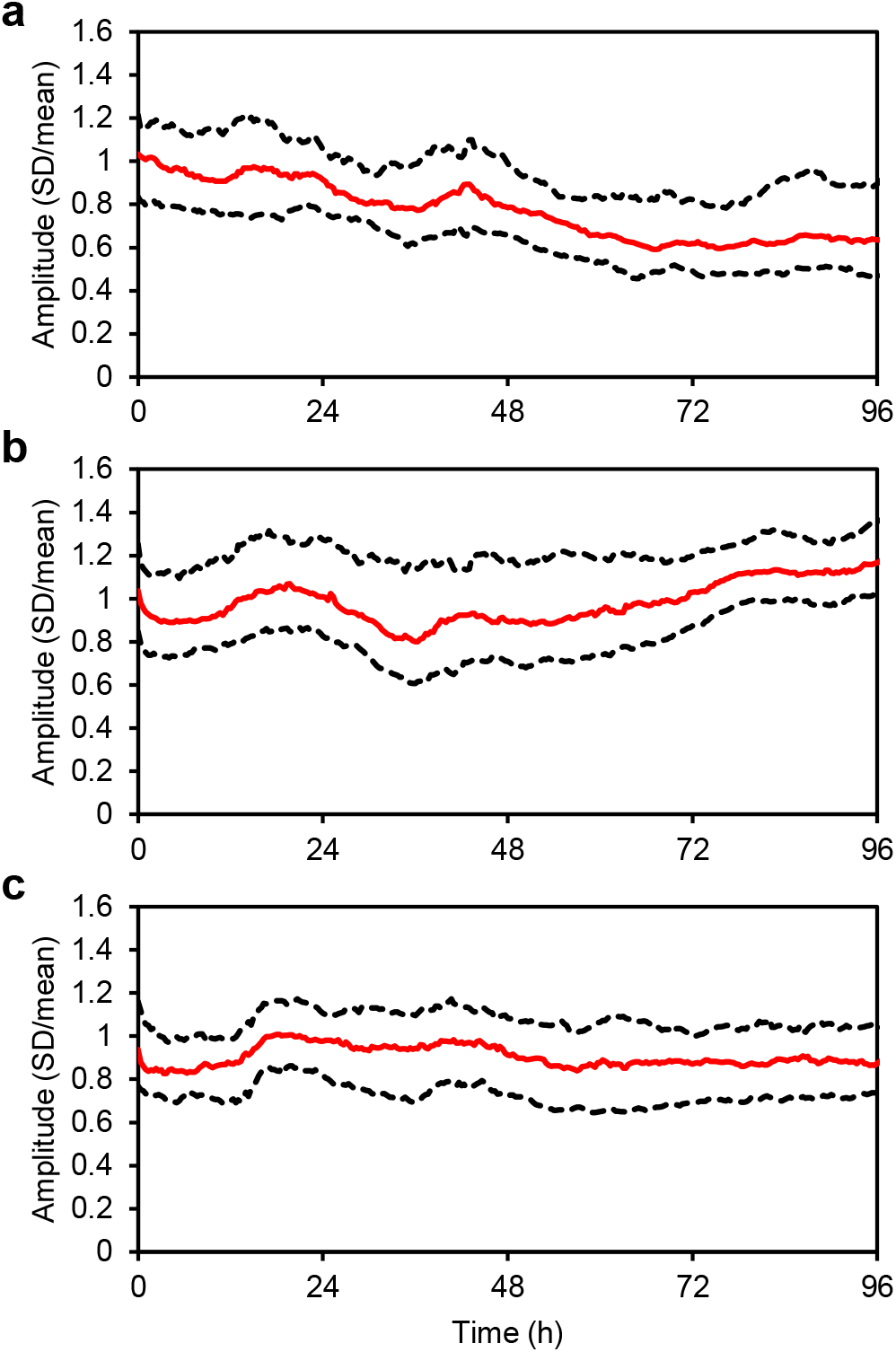
Temporal changes in the amplitudes of circadian rhythms of leaf-derived cells Amplitude of a cellular rhythm at a particular time was defined as the coefficient of variation (standard deviation/mean) of bioluminescence intensities in the following 24 h range with 72 time points. Cellular rhythms of cells with circadian rhythm under standard- (**a**; 80 leaf-derived cells) and high-cell-density conditions (**b**; 153 leaf-derived cells), and in conditioned medium (**c**; 109 leaf-derived cells) were analyzed. A red line represents medium and dashed lines represent 25% and 75% quartiles in each graph.

**Supplementary Table 1.** Summary of the periods of individual cells with circadian rhythm under standard conditions

| Exp. index | Cell types | Cells with circadian rhythm |  |  |
| --- | --- | --- | --- | --- |
| | | FRP ±SD (h) | Cellular periods ±SD (h) | CV of peak intervals |
| Exp1 | Leaf-derived cells | 22.9 ± 2.3 | 22.1 ± 1.7 | 0.15 |
| Exp2 | Leaf-derived cells | 23.5 ± 3.8 | 23.5 ± 2.3 | 0.16 |
| Exp3 | Leaf-derived cells | 25.9 ± 1.7 | 22.2 ± 1.8 | 0.27 |
| Exp4 | Root-derived cells | 25.0 ± 2.9 | 24.8 ± 2.0 | 0.08 |
| Exp5 | Root-derived cells | 23.1 ± 2.6 | 23.6 ± 1.5 | 0.09 |

**Supplementary Table 2.** Summary of the ratio of individual cells with circadian-rhythm in a high-cell-density culture in three independent experiments

| Exp. Index | Number of cells (percentages) |  |  |
| --- | --- | --- | --- |
|  | Group 0 | Group 1 | Group 2 |
| Exp33 | 66 (47.1) | 10 (7.1) | 64 (45.7) |
| Exp34 | 51 (42.5) | 12 (10.0) | 57 (47.5) |
| Exp35 | 36 (27.5) | 21 (16.0) | 74 (56.5) |
| Total | 153 (39.1) | 43 (11.0) | 195 (49.9) |

## REFFERENCES

1. Pittendrigh, C. S. Circadian rhythms and the circadian organization of living systems. Cold Spring Harbor Symposia on Quantitative Biology 25, 159–184 (1960).

2. Roenneberg, T. & Deng, T. S. Photobiology of the Gonyaulax circadian system. I. Different phase response curves for red and blue light. Planta 202, 494–501 (1997).

3. Barak, S., Tobin, E. M., Green, R. M., Andronis, C. & Sugano, S. All in good time: the Arabidopsis circadian clock. Trends in Plant Science 5, 517–522 (2000).

4. Millar, A. J. A suite of photoreceptors entrains the plant circadian clock. Journal of Biological Rhythms 18, 217–226 (2003).

5. Creux, N. & Harmer, S. Circadian rhythms in plants. Cold Spring Harbor Perspectives in Biology 11, 1–18 (2019).

6. Mcclung, R. The plant circadian oscillator. biology 8, 1–17 (2019).

7. Pittayakanchit, W., Lu, Z., Chew, J., Rust, M. J. & Murugan, A. Biophysical clocks face a trade-off between internal and external noise resistance. eLife 7, 1–38 (2018).

8. Gonze, D., Halloy, J. & Goldbeter, A. Robustness of circadian rhythms with respect to molecular noise. Proceedings of the National Academy of Sciences of the United States of America 99, 673–678 (2002).

9. Cortijo, S., Aydin, Z., Ahnert, S. & Locke, J. C. Widespread inter-individual gene expression variability in Arabidopsis thaliana. Molecular systems biology 15, e8591 (2019).

10. James, A. B. et al. The circadian clock in Arabidopsis roots is a simplified slave version of the clock in shoots. Science 322, 1832–1835 (2008).

11. Bordage, S., Sullivan, S., Laird, J., Millar, A. J. & Nimmo, H. G. Organ specificity in the plant circadian system is explained by different light inputs to the shoot and root clocks. The New phytologist 212, 136–149 (2016).

12. Takahashi, N., Hirata, Y., Aihara, K. & Mas, P. A Hierarchical multi-oscillator network orchestrates the Arabidopsis circadian system. Cell 163, 148–159 (2015).

13. Gould, P. D. et al. Coordination of robust single cell rhythms in the Arabidopsis circadian clock via spatial waves of gene expression. eLife 7, e31700 (2018).

14. Nohales, M. A. & Kay, S. A. Molecular mechanisms at the core of the plant circadian oscillator. Nature Structural and Molecular Biology 23, 1061–1069 (2016).

15. Sorkin, M. L. & Nusinow, D. A. Time Will Tell: Intercellular communication in the plant clock. Trends in plant science (2021).

16. Yakir, E. et al. Cell autonomous and cell-type specific circadian rhythms in Arabidopsis. Plant Journal 68, 520–531 (2011).

17. Okada, M., Muranaka, T., Ito, S. & Oyama, T. Synchrony of plant cellular circadian clocks with heterogeneous properties under light/dark cycles. Scientific Reports 7, 1–10 (2017).

18. Kanesaka, Y., Okada, M., Ito, S. & Oyama, T. Monitoring single-cell bioluminescence of Arabidopsis leaves to quantitatively evaluate the efficiency of a transiently introduced CRISPR/Cas9 system targeting the circadian clock gene ELF3. Plant biotechnology 36, 187–193 (2019).

19. Watanabe, E., Isoda, M., Muranaka, T., Ito, S. & Oyama, T. Detection of uncoupled circadian rhythms in individual cells of Lemna minor using a dual-color bioluminescence monitoring system. Plant and Cell Physiology (2021).

20. Muranaka, T. & Oyama, T. Heterogeneity of cellular circadian clocks in intact plants and its correction under light-dark cycles. Science advances 2, e1600500 (2016).

21. Fukuda, H., Ukai, K. & Oyama, T. Self-arrangement of cellular circadian rhythms through phase-resetting in plant roots. Physical Review E Statistical, Nonlinear, and Soft Matter Physics 86, 1–5 (2012).

22. Fukuda, H., Nakamichi, N., Hisatsune, M., Murase, H. & Mizuno, T. Synchronization of plant circadian oscillators with a phase delay effect of the vein network. Physical Review Letters 99, 1–4 (2007).

23. Wenden, B., Toner, D. L. K., Hodge, S. K., Grima, R. & Millar, A. J. Spontaneous spatiotemporal waves of gene expression from biological clocks in the leaf. Proceedings of the National Academy of Sciences 109, 6757–6762 (2012).

24. Greenwood, M., Domijan, M., Gould, P. D., Hall, A. J. W. & Locke, J. C. W. Coordinated circadian timing through the integration of local inputs in Arabidopsis thaliana. PLoS biology 17(8), e3000407 (2019).

25. Kim, J. & Somers, D. E. Rapid assessment of gene function in the circadian clock using artificial microRNA in Arabidopsis mesophyll protoplasts. Plant Physiology 154, 611–621 (2010).

26. Nakamichi, N., Matsushika, A., Yamashino, T. & Mizuno, T. Cell autonomous circadian waves of the APRR1/TOC1 quintet in an established cell line of Arabidopsis thaliana. 44, 360–365 (2003).

27. Nakamura, S. & Oyama, T. Long-term monitoring of bioluminescence circadian rhythms of cells in a transgenic Arabidopsis mesophyll protoplast culture. Plant biotechnology 35, 291–295 (2018).

28. Nakamichi, N. et al. Characterization of plant circadian rhythms by employing Arabidopsis cultured cells with bioluminescence reporters. Plant & cell physiology 45, 57–67 (2004).

29. Nakamichi, N., Kita, M., Ito, S., Yamashino, T. & Mizuno, T. PSEUDO-RESPONSE REGULATORS, PRR9, PRR7 and PRR5, together play essential roles close to the circadian clock of Arabidopsis thaliana. Plant & cell physiology 46, 686–98 (2005).

30. Yoo, S. D., Cho, Y. H. & Sheen, J. Arabidopsis mesophyll protoplasts: A versatile cell system for transient gene expression analysis. Nature Protocols 2, 1565–1572 (2007).

31. Petersson, S. v. et al. An auxin gradient and maximum in the Arabidopsis root apex shown by high-resolution cell-specific analysis of IAA distribution and synthesis. The Plant cell 21, 1659–68 (2009).

32. Pasternak, T., Paponov, I. A. & Kondratenko, S. Optimizing protocols for Arabidopsis shoot and root protoplast cultivation. Plants 10, 1–17 (2021).

33. Johnson, C. H. Phase Response Curves: What can they tell us about circadian clocks. Circadian Clocks from Cell to Human 209–246 (1992).

34. Schmal, C., Herzog, E. D. & Herzel, H. Measuring relative coupling strength in circadian systems. Journal of Biological Rhythms 33, 84–98 (2018).

35. Dalchau, N. et al. The circadian oscillator gene GIGANTEA mediates a long-term response of the Arabidopsis thaliana circadian clock to sucrose. Proceedings of the National Academy of Sciences of the United States of America 108, 5104–5109 (2011).

36. Bieniawska, Z. et al. Disruption of the Arabidopsis circadian clock is responsible for extensive variation in the cold-responsive transcriptome. Plant Physiology 147, 263–279 (2008).

37. Gould, P. D. et al. The molecular basis of temperature compensation in the Arabidopsis circadian clock. Plant Cell 18, 1177–1187 (2006).

38. Endo, M., Shimizu, H., Nohales, M. A., Araki, T. & Kay, S. A. Tissue-specific clocks in Arabidopsis show asymmetric coupling. Nature 515, 419–422 (2014).

39. Illston, B. G. & Fiebrich, C. A. Horizontal and vertical variability of observed soil temperatures. Geoscience Data Journal 4, 40–46 (2017).

40. Nimmo, H. G. Entrainment of Arabidopsis roots to the light:dark cycle by light piping. Plant Cell and Environment 41, 1–7 (2018).

41. Johnson, C. H., Elliott, J. A. & Foster, R. Entrainment of circadian programs. Chronobiology International 20, 741–774 (2003).

42. Murashige, T. & Skoog, F. A Revised medium for rapid growth and bioassays with tobacco tissue cultures. Physiologia Plantarum 15, 473–497 (1962).

43. Muranaka, T. & Oyama, T. Application of single-cell bioluminescent imaging to monitor circadian rhythms of individual plant cells. in Methods in Molecular Biology vol. 2081 231–242 (Humana Press Inc., 2020).

44. Isoda, M. & Oyama, T. Use of a duckweed species, Wolffiella hyalina, for whole-plant observation of physiological behavior at the single-cell level. Plant Biotechnology 35, 387–391 (2018).

45. Muranaka, T., Okada, M., Yomo, J., Kubota, S. & Oyama, T. Characterisation of circadian rhythms of various duckweeds. Plant Biology 17, 66–74 (2015).

